# Dynamic *Ins2* gene activity defines β-cell maturity states

**DOI:** 10.1101/702589

**Authors:** Chieh Min Jamie Chu, Honey Modi, Søs Skovsø, Cara Ellis, Nicole A.J. Krentz, Yiwei Bernie Zhao, Haoning Cen, N Noursadeghi, Evgeniy Panzhinskiy, Xiaoke Hu, Derek A. Dionne, Yi Han Xia, Shouhong Xuan, Mark O. Huising, Timothy J. Kieffer, Francis C. Lynn, James D. Johnson

## Abstract

Transcriptional and functional cellular specialization has been described for insulin-secreting β-cells of the endocrine pancreas. However, it is not clear whether β-cell heterogeneity is stable or reflects dynamic cellular states. We investigated the temporal kinetics of endogenous insulin gene activity using live cell imaging, with complementary experiments employing FACS and single cell RNA sequencing, in β-cells from *Ins2*^GFP^ knock-in mice. *In vivo* staining and FACS analysis of islets from *Ins2*^GFP^ mice confirmed that at a given moment, ~25% of β-cells exhibited significantly higher activity at the conserved insulin gene *Ins2.* Live cell imaging captured *Ins2* gene activity dynamics in single β-cells over days. Autocorrelation analysis revealed a subset of cells with oscillating behavior, with mean oscillation periods of 17 hours. Increased glucose concentrations stimulated more cells to oscillate and resulted in higher average *Ins2* gene activity per cell. Single cell RNA sequencing showed that *Ins2*(GFP)^HIGH^ β-cells were enriched for markers of β-cell maturity. *Ins2*(GFP)^HIGH^ β-cells were also significantly less viable at all glucose concentrations and in the context of ER stress. Collectively, our results demonstrate that the heterogeneity of insulin production, observed in mouse and human β-cells, can be accounted for by dynamic states of insulin gene activity.

**Blurb:** Previously reported pancreatic β-cell heterogeneity reflects β-cell state transitions.

## Introduction

Pancreatic β-cells in the islets of Langerhans are the only source of circulating insulin, a conserved and essential hormone that is required for nutrient homeostasis and life (1). Insulin production is demanding, as insulin mRNA can account for roughly half of all β-cell mRNA and its synthesis, folding and processing require semi-specialized transcription factors, enzymes and cellular conditions (2; 3). However, not all β-cells appear the same. Indeed, functional β-cell heterogeneity is well established (4), including cellular specialization for islet cell synchronization, insulin secretion, insulin production, and marker gene expression (5–11). For example, recent *in situ* imaging has revealed the existence of extreme β-cells, defined as having >2-fold *Ins2* mRNA than the median expression as measured by single-molecule fluorescence *in situ* hybridization (4; 12). Single cell RNA sequencing has shown that human β-cells also express *INS* over a similarly wide range (8). However, it remains unclear whether this variation is the hallmark of distinct stable populations of β-cells, or whether it is indicative of transitions between more labile β-cell states.

To date, the vast majority of islet cell ‘sub-populations’ have been defined by single time-point snapshots, making it impossible to know to what extent observed β-cell heterogeneity represents distinct islet cell ‘fates’ or variations in islet cell ‘states’. This information is essential for interpreting existing and future data in the field. Using a dual *Ins1* and *Pdx1* promoter reporter construct and live cell imaging, we have previously demonstrated that mouse and human β-cells can transition between less and more differentiated states over a time scale of ~24 hours (13–15). However, the artificial promoter constructs in these early studies may not reflect endogenous gene activity, leaving open the question of whether endogenous insulin gene activity is similarly dynamic.

In this study, we measured endogenous insulin gene activity using an *Ins2*^GFP^ knock-in/knockout mouse line in which the coding sequencing of the evolutionarily conserved *Ins2* gene has been replaced with GFP (16). Live cell imaging of dispersed cells from these mice revealed that GFP fluorescence changed over time in a sub-set of cells, suggesting that variation in *Ins2* levels results from dynamic transcriptional activity at the *Ins2* gene locus rather than stable heterogeneity. Single cell RNA sequencing was used to characterize the *Ins2*(GFP)^HIGH^ cellular state in an unbiased way, revealing increased markers of β-cell maturity as well as alterations in protein synthesis machinery and cellular stress response networks. Pancreatic β-cells in the *Ins2*(GFP)^HIGH^ cellular state were also more fragile across a range of stress conditions. To the best of our knowledge, our observations are the first to define the temporal kinetics of endogenous insulin gene activity, which represents a previously uncharacterized form of β-cell plasticity. Understanding the dynamics of insulin production has relevance for understanding the pathobiology of diabetes and for regenerative therapy research (17).

## Methods

### Animals and in vivo physiology

Animals were housed and studied in the Modified Barrier Facility using protocols approved by the UBC Animal Care Committee in accordance with international guidelines. *Ins2*^WT/GFP^ knock-in mice were obtained from Shouhong Xuan (16). These mice were crossed with transgenic mice where the *Ins1* promoter drives a mCherry:H2B fluorescent fusion protein (18). Glucose tolerance and insulin secretion were assessed in both male and female mice (12-14 weeks) injected intraperitoneally with 2g/kg (20%) glucose after a 5 hr fast. Insulin from *in vivo* samples was measured using ELISA kits from Alpco (Salem, NH, USA). Insulin tolerance was assessed after injection of 0.75 U insulin per kg body weight after a 5 hr fast.

### Immunostaining and RT-PCR

Pancreata from PBS perfused mice were harvested and fixed in 4% paraformaldehyde for 24 hr before being washed and stored in 70% ethanol, prior to paraffin embedding. Pancreatic sections (5 μm) were taken from at least three different regions of the pancreas 100 μm apart. Sections were deparaffinized, hydrated with decreasing concentrations of ethanol, and rinsed with PBS. Sections were subjected to 15 min of heat-induced epitope retrieval at 95°C using a 10 mM citrate buffer, pH 6.0. Sections were blocked then incubated with primary antibodies overnight in a humid chamber at 4°C. A list of primary antibodies can be found in Antibodies section and Table S1. Primary antibodies were visualized following incubation with secondary antibodies conjugated to AlexaFluor 488, 555, 594, or 647 as required (1:1,000; Invitrogen). Counter staining was done by Vectasheild mounting media with DAPI (H-1200). Images for β-cell and α-cell area were taken on ImageXpress^MICRO^ using a 10× (NA 0.3) objective and analyzed using the MetaXpress software (Molecular Devices Corporation, San Jose, CA, USA). All other images were taken on a Zeiss 200M microscope using 20× air (NA 0.75), 40× oil (NA 1.3), and/or 100× oil (NA 1.45) objectives and analyzed using Slidebook software (Intelligent Imaging Innovations, Denver, CO, USA). For quantification of immunofluorescence, we used the segment masking function of Slidebook, generating a GFP high mask and a GFP low mask and comparing the two masks (Fig. 8C). For human pancreas sections stained for C-peptide and PDX1, we obtained images from the HPAP database (19) and performed analysis with custom scripts in CellProfiler. Real-time RT-PCR was conducted as described previously (1). A list of primers used can be found in Table S2.

### Islet isolation, dissociation and culture

Pancreatic islets were isolated using collagenase, filtered, and hand-picked as previously described (15). Islets were cultured overnight (37°C, 5% CO_2_) in RPMI1640 medium (Thermofisher) with 11 mM glucose (Sigma), 100 units/ml penicillin, 100 μg/ml streptomycin (Thermofisher), and 10% vol/vol FBS (Thermofisher). For islet dissociation in preparation for imaging experiments, islets were washed 4 times with MEM (Corning) and immersed for 5 minutes in 0.05% trypsin at 37°C. Cells were then resuspended in RPMI1640 medium and given 24 hours of rest (37°C, 5% CO_2_) before imaging. For experiments involving various culture conditions, cells were incubated overnight in serum-free media for serum-free and serum-starved conditions, and media with 10% FBS for serum conditions. Prior to imaging, cells were given a media change and stained with Hoechst cell nuclei marker and propidium iodide cell death marker for 2 hours. Serum-free condition cells were given media with no serum, while serum-starved (rescue) and serum conditions were given media with 10% FBS. Cells were then treated with various glucose concentrations immediately prior to imaging.

### Fluorescence-activated cell sorting

Pancreatic islets were dispersed using 0.05% trypsin and resuspended in 1xPBS with 0.005% FBS. Dispersed islets were then filtered into 5 ml polypropylene tubes. FACS was conducted on a Cytopeia Influx sorter (Becton Dickinson, Franklin Lakes, NJ, USA) at the Life Sciences Institute core facility. Cells were excited with a 488 nm laser (530/40 emission) and a 561 □nm laser (610/20 emission).

### Live cell imaging

To define the incidence and kinetics of the transitions in *Ins2* gene activity, we cultured dispersed islet cells on 96-well glass bottom plates, and imaged them every 5 or 30 minutes for up to 96 hours through a 40x air objective using a ImageXpress^MICRO^ environmentally-controlled, robotic imaging system (Molecular Devices) employing a 300 W Xenon lamp (20). Cells were exposed to 359 nm light for 110 ms, 491 nm light for 15 ms, 561 nm for 75 ms.

Our methods for live cell imaging of islet cell survival have been published (20). Briefly, pancreatic islet cells were dissociated, and cultured in RPMI1640 media with 10% FBS (Gibco, Fisher Scientific, Gaithersburg, MD, USA), and penicillin/streptomycin on 96 well glass bottom plates for 48 hours. Dispersed islet cells were then exposed to different glucose concentrations, as well as thapsigargin (Sigma, St. Louis, MO, USA). Islet cell death was measured by propidium iodide incorporation. Images were taken every 30 min for up to 48 hours, as described earlier. Propidium iodide incorporation was traced throughout the time period, and area under the curve was measured.

### Live-cell imaging analysis

Analysis of *Ins2*^GFP^:*Ins1*-mCherry cells was done using ImageXpress software and custom R scripts (21). Hierarchical clustering with 8 cellular behavior traits as features was done, and a heatmap was constructed using the pheatmap function. Traits were calculated using R functions and formulas, including mean (average life time fluorscent intensity), sharp (calculates number of sharp peaks in a cell’s lifetime), AUC (area under curve), FWHM (full width at half maximum), and oscillation (mean average deviation). Principle component analysis was done using the same 8 cellular behavior traits as variables. We performed model-based clustering using the mclust function in R, and projected the cells onto the first two principle components.

Analysis of *Ins2*^GFP^ cells was done using MetaXpress analysis software and custom R scripts. Hierarchical clustering was done with various cellular behavior traits as features, and a heatmap was constructed using the pheatmap function in R. Features include ACF time (calculates when the autocorrelation function equals 0 in a time series), FWHM (full width at half maximum), difference (calculates the range of fluorescent intensity change), deviation (mean average deviation), mean (average lifetime relative fluorescent intensity), and sharp (a function which detects sharp peaks). Principle component analysis was done using the same cellular behavior traits as variables. We performed K-means clustering using the kmeans function in R, and projected the cells onto the first two principle components. Categorization of cells into high and low GFP expressing states in live cell imaging and single cell RNA sequencing data was done using model based clustering (22).

We noticed in our imaging experiments a slow decrease in GFP fluorescence over the course of 48 hours (Supplemental Figure 4A). An experiment with 2-hour imaging intervals and fewer wells showed that decreasing the total exposure time over 48 hours of imaging did not remove this trend (Supplemental Figure 4B). De-trending and normalizing to control for this trend was done by comparing various treatments to the basal (10 mM glucose and normalizing to the average fluorescence intensity of first 2 hours of imaging to correct for a possible linear artifact and variations in the initial hours of imaging, using the following calculation:

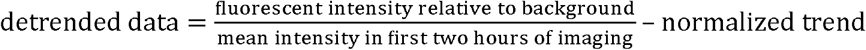

Autocorrelation analysis of single mutant *Ins2*^WT/GFP^ was done using the acf function in R (23). To calculate the oscillation periods of GFP, we first selected for cells with at least one oscillation. We calculated the autocorrelation function of all cells for all time-points, and filtered for cells with at least two significantly positive peaks (autocorrelation function >0.05) using a find peaks function in R (21), indicating an oscillation. We calculated oscillation periods by finding the distance of the two positive peaks to get a full wavelength period. Oscillation periods of oscillating cells with more than one oscillation were then binned in 4-hour intervals and plotted into a barchart using R scripts.

### Single cell transcriptomics

Islets were isolated, hand-picked, dispersed, and subjected to FACS as described above into *Ins2*(GFP)^HIGH^ and *Ins2*(GFP)^LOW^ cells. Single Cell Suspension was loaded on the 10x Genomics Chromium Controller for capture in droplet emulsion. The 60-week old mouse islet cell libraries were prepared using the Chromium Single Cell 3’ Reagent v2 Chemistry kit (10x Genomics, Pleasanton, CA, USA) and the standard protocol was followed for all steps. The 8-week old mouse islet cell libraries were prepared using the Chromium Single Cell 3’ Reagent v3 Chemistry kit (10x Genomics). Libraries were then sequenced on a Nextseq500 (Illumina). Cell Ranger 3.0 (10x Genomics) was used to process raw fastq data. The results were visualized using 10x Genomics Loupe Cell Browser. Cells with high mitochondria gene ratios (>0.1), low numbers of genes (<500), and low UMI counts (<1000) were excluded prior to analysis. This was done for both 60-week-old mouse data and 8-week-old mouse data respectively.

Seurat R package (version 4.0.4) was used to perform analysis. (24). Data were normalized, scaled and analyzed with PCA. PCs generated were used to build a UMAP, with which we used to identify populations of cells using known cellular markers (*Ins1* for β-cells, *Gcg* for α-cells, *Sst* for δ cells, *Ppy* for pancreatic polypeptide/ γ cells, *Ghrl* for ε cells, and *Pcam1* for endothelial cells). Cells expressing *Ins1* were selected, and the rest were used as a base expression level for differential expression analysis using the *FindMarkers* function in Seurat. Differential expression analysis between the subsets of cells was performed using the Wilcoxon rank sum test option in Seurat. Differential expression analysis over GFP gradient was performed by first calculating correlation functions for each gene with GFP, then selecting for the genes that had the highest values. Top hits were used for pathway enrichment analyses and heatmaps. Pathway enrichment analysis was performed using custom R scripts with data from Gene Ontology Resource database. RNA velocity analysis was performed by mapping reads to exons and introns with velocyto, while RNA velocity vectors were inferred and overlaid onto UMAPs using ‘dynamical’ mode in scVelo.

### Statistical analysis

Data are shown as mean ± SEM unless otherwise indicated. Differences between 2 groups were evaluated with Student’s t-test and between more than 2 groups using ANOVA and Kruskal–Wallis one-way analysis of variance, and were calculated with custom R-scripts or Prism software (Graphpad Software, San Diego, CA, USA).

### Data resource and availability

The datasets generated during and/or analyzed during the current study are available from the corresponding author upon reasonable request. Key resources are stated in Table 1. Additional info may be found in Table S1.

**Table 1.**
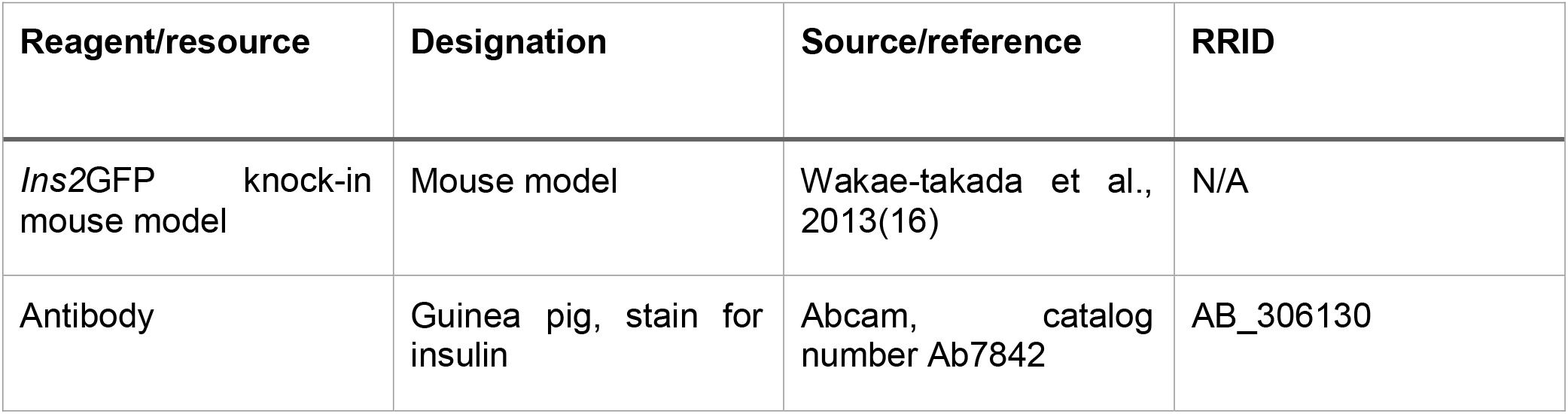
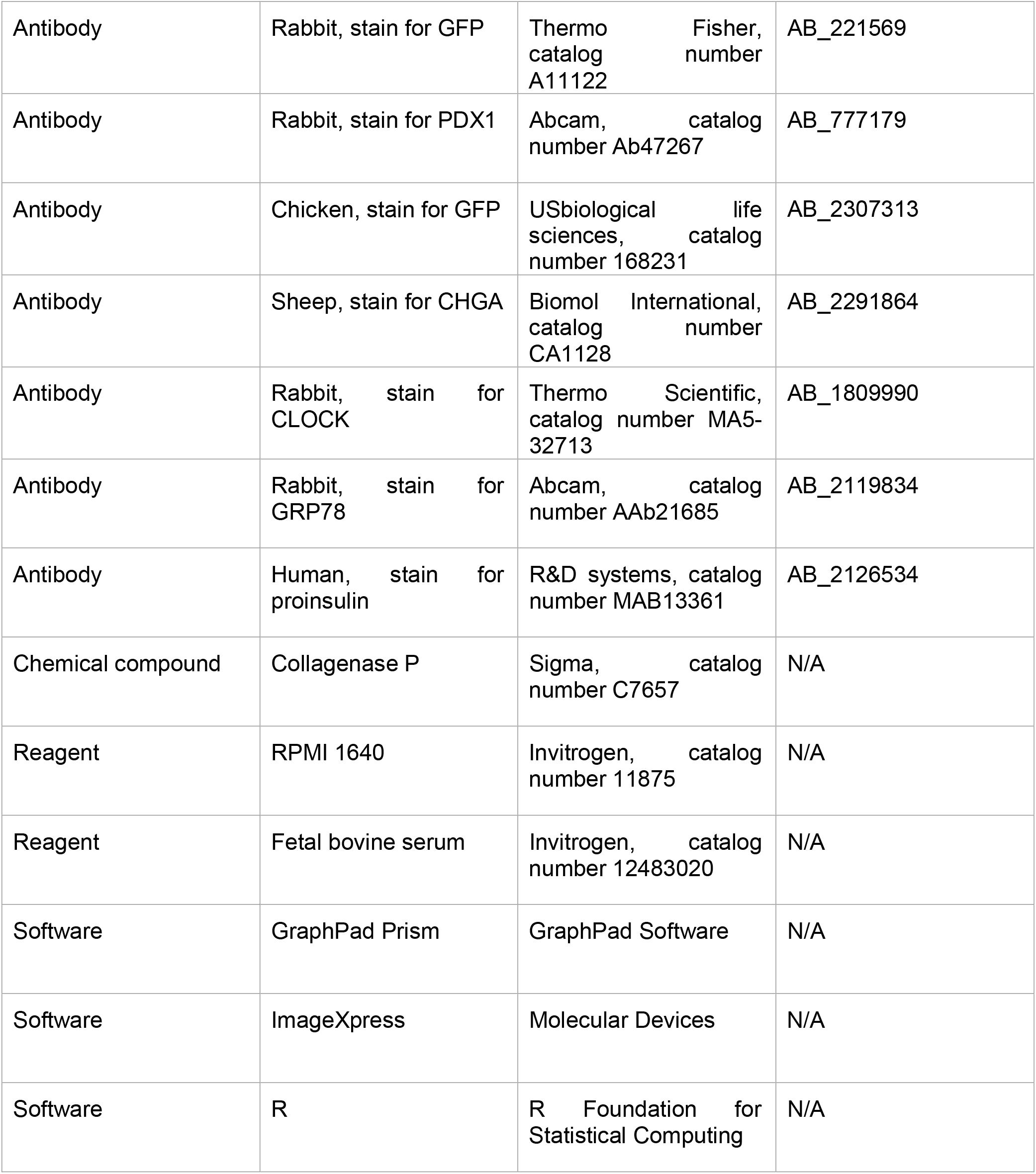
Key resources.

## Results

### In vivo heterogeneity of insulin content in human β-cell and insulin gene activity in Ins2^GFP^ mice

Heterogeneity of insulin production is an established phenomenon. We started our study by investigating human pancreases from online databases (19) that were stained with antibodies to C-peptide and PDX1, a key transcription factor for β-cell survival and function (25; 26). As expected, based on single cell RNA sequencing data (8) and previous single cell imaging of human β-cells (27),we identified β-cells with both high and low C-peptide and PDX1 protein levels in human pancreas (Fig. 1A, B, C). We have previously shown limited correlations between insulin content and PDX1 immunofluorescence or nuclear localization in human β-cells (27). Analysis of single cell RNA sequencing data previously compiled by our lab (2) showed that there was a bimodal distribution in insulin gene expression in β-cells from both type 2 diabetic and non-diabetic donors, with a left-shift in high insulin expressing cells in patients with type 2 diabetes (Fig 1D).

**Figure 1.**
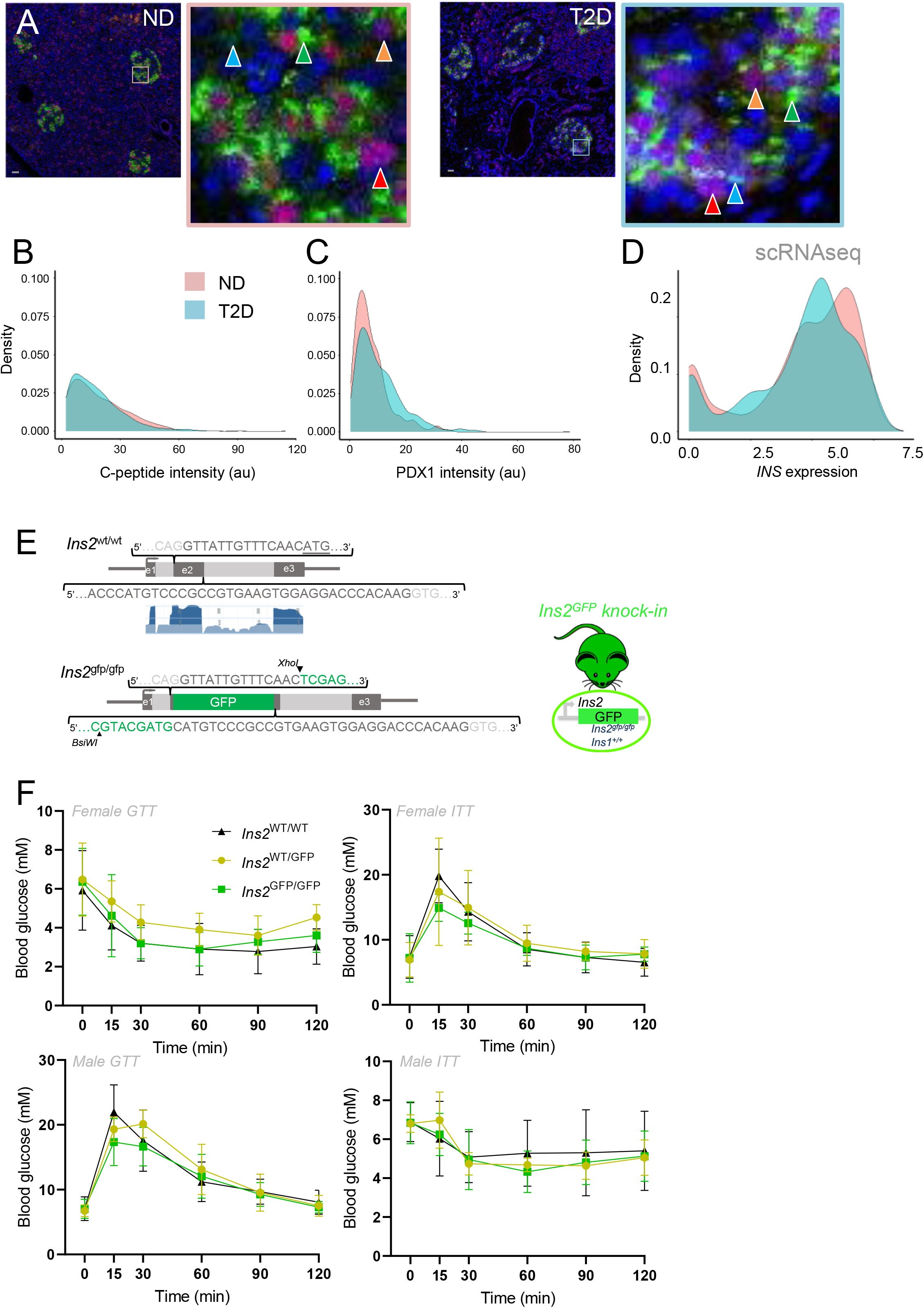
*In vivo* heterogeneity of insulin content and gene activity in human and mouse islets. **(A)** Variability in C-peptide (green stain) and PDX1 (red stain) protein levels in pancreatic sections from non-diabetic and type-2 diabetic donors. Orange arrows indicate low PDX1 expressing cells, red arrows indicate high PDX1 expressing cells, blue arrows indicate low C-peptide expressing cells, and green arrows indicate high C-peptide expressing cells. Scale bar is 100 μm. **(B, C)** Quantification of staining for C-peptide and PDX1 in pancreatic sections from non-diabetic and type-2 diabetic donors, displayed as a histogram. **(D)** Comparison of insulin mRNA distribution in non-diabetic and type-2 diabetic human β-cell single cell RNA sequencing data. **(E)**The *Ins2*^GFP/GFP^ knock-in mouse model. Illustration shows, with scale, the replacement of most of the second exon with wildtype GFP. The wildtype *Ins2* locus is shown for comparison, including RNA-seq exon coverage, aggregate (filtered) from NCBI *Mus musculus* Annotation release 106 (log 2 base scaled). **(F)** Normal glucose homeostasis in *Ins2*^WT/GFP^, *Ins2*^GFP/GFP^ mice and WT (Male n=4-8, Female n=4-5).

In order to study insulin gene activity in living cells, we examined islets from mice in which the coding sequencing of the evolutionarily conserved *Ins2* gene has been replaced with GFP (16) (Fig. 1E). Mice lacking 1 or 2 wildtype functional *Ins2* alleles had normal glucose homeostasis (Fig. 1F), consistent with our previous studies of *Ins2* knockout mice (28) and the ability of *Ins1* to compensate (29). Immunofluorescence staining of pancreata from *Ins2*^GFP^ mice revealed a possible bimodal distribution of endogenous insulin production *in vivo,* similar to our single cell analyses (Fig. 2A). Hand counting 1879 cells across 11 randomly selected islets showed that 38.7% of β-cells had substantially higher GFP immunofluorescence above an arbitrary, but consistently applied, threshold (Fig. 2A). The percentage of cells with high GFP did not appear to vary as a function of islet size. We have observed similar heterogeneity when examining the β-galactosidase knock-in in to the *Ins2* locus (28), meaning that this observation is not unique to this knock-in line or to GFP as a reporter.

**Figure 2.**
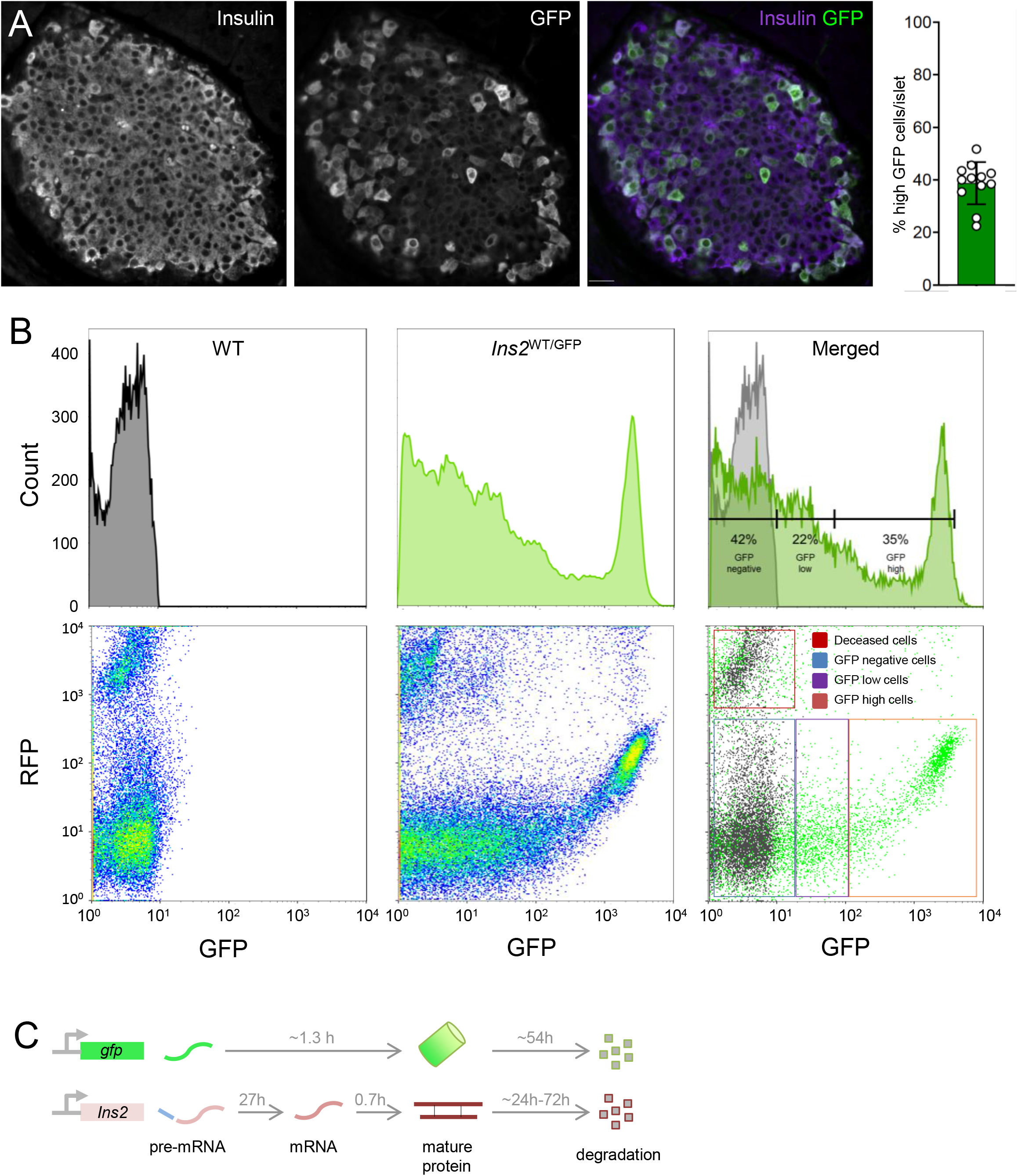
Identifying high and low GFP expressing cells in *Ins2*^WT/GFP^ knock-in mice. **(A)** GFP from the endogenous *Ins2* gene locus is high in a subset of mouse pancreatic β-cells. Scale bar is 20 μm. **(B)** Detection of *Ins2*(GFP)^LOW^ and *Ins2*(GFP)^HIGH^ populations of β-cells by FACS. Includes β-cells isolated from a WT mouse for comparison. **(C)** Presumed transcription-to-maturation times of GFP and insulin. See Supplemental Information for calculations and references.

FACS confirmed this bimodal distribution and that less than half of all β-cells engage in high *Ins2* gene transcription at a given time (Fig. 2B). FACS analysis also validated that *gfp* mRNA, *Ins2* mRNA, and pre-mRNA were significantly increased in high GFP-expressing cells compared to cells expressing less GFP (Supplemental Fig. S1A), strongly suggesting that GFP protein levels accurately reflect *Ins2* mRNA in this system. We did not expect there to be a perfect correlation due to the different predicted transcription-to-protein time courses for GFP and insulin (Fig. 2C; see Supplemental Information for calculations). Nevertheless, these data demonstrate that GFP production, reflecting the activity of the endogenous *Ins2* gene locus, is bimodal *in vivo* and *ex vivo,* with 35% of β-cells showing significantly higher activity. Hereafter, we refer to cells with high GFP abundance as *Ins2*(GFP)^HIGH^.

### Live cell imaging of insulin gene activity in Ins2^WT/GFP^ mice

We next generated a mouse model to examine endogenous *Ins2* activity dynamics relative to a stable background β-cell marker. We crossed the *Ins2*^GFP^ knock-in line with transgenic mice with an allele wherein histone-fused mCherry is driven by the less complex *Ins1* promoter that is known to have relatively stable red fluorescence in virtually all β-cells (18), allowing us to track *Ins2* gene activity in real-time (GFP), while observing all β-cells (mCherry). As expected, immunofluorescence of intact pancreatic sections and FACS analysis of dispersed islets from *Ins2*^WT/GFP^:*Ins1*-mCherry mice showed that mCherry labelled virtually all β-cells, while GFP was robustly expressed in a clearly separated sub-set of β-cells we deemed *Ins2*(GFP)^HIGH^ (Fig. 3A,B,C). qPCR of these FACS purified cells confirmed the expected elevated expression of GFP, *Ins2, pre-Ins2, Ins1,* and pre-*Ins1* mRNA in *Ins2*(GFP)^HIGH^ cells (Fig. 3D, Supplemental Fig. S1B). *Ins2* pre-mRNA would be expected to precede GFP fluorescence by at least 1.3 hours, as per estimations in Fig. 2C and calculations in Supplemental Information, which provides a possible explanation for the elevated pre-*Ins2* in the mCherry positive, but GFP low β-cells. Both intact islets and dispersed islet cells isolated from *Ins2*^WT/GFP^:*Ins1*-mCherry mice showed a similar proportion of *Ins2*(GFP)^HIGH^ and *Ins2*(GFP)^LOW^ cells to that we observed *in vivo,* demonstrating that this heterogeneity was not altered by isolation or dispersion/culture.

**Figure 3.**
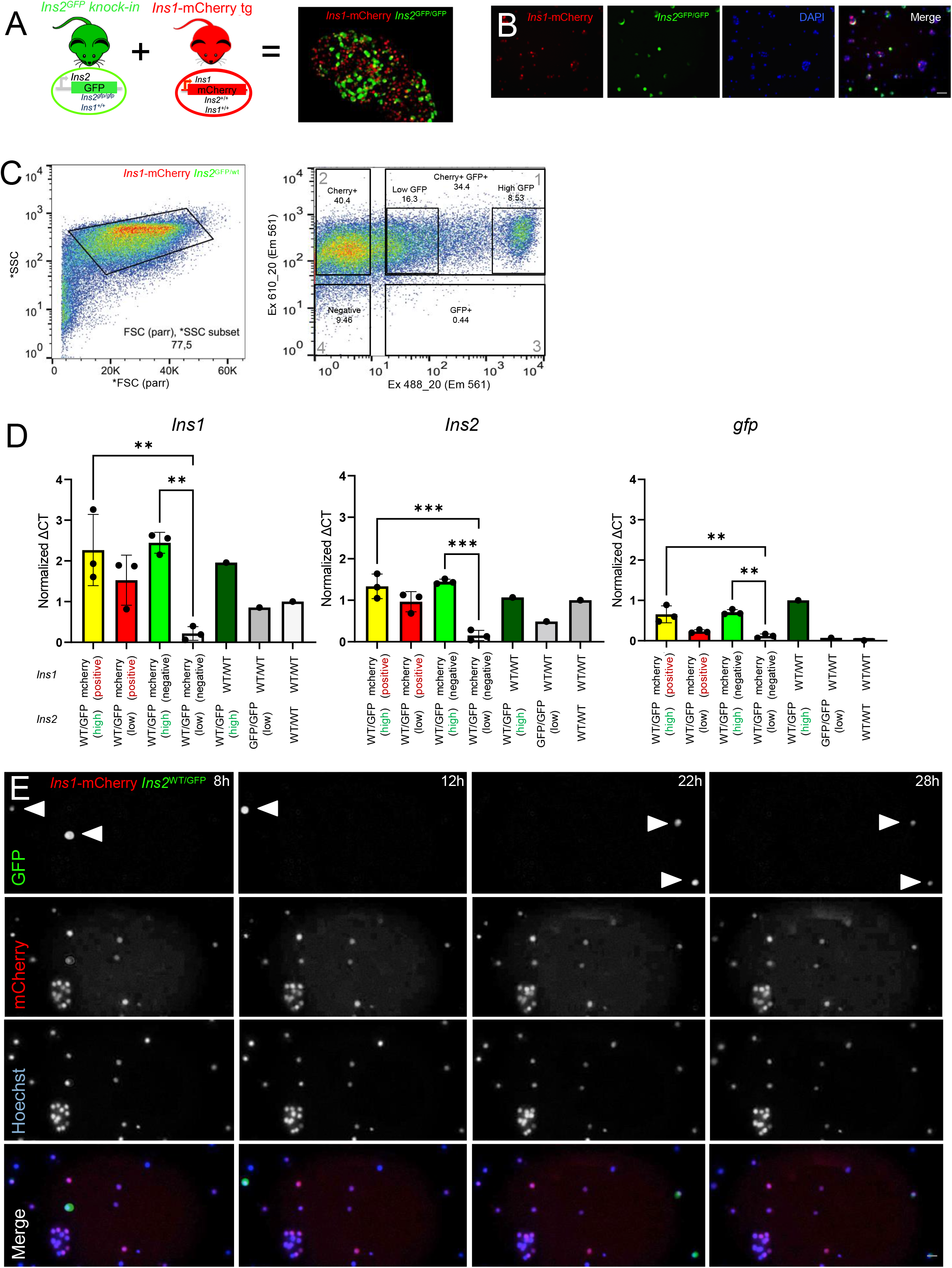
Tracking β-cell heterogeneity with *Ins1*-mCherry;*Ins2*^WT/GFP^ knock-in mice. **(A)** Experimental model for tracking activity at the endogenous *Ins2* locus (GFP) in β-cells marked with mCherry driven by an *Ins1* promoter, which constitutively marks 98% of β-cells. Scale bar is 50 μm. **(B)** Live cell imaging of islet cells isolated and dispersed from *Ins1*-mCherry;*Ins2*^WT/GFP^ mice. Scale bar is 50 μm. **(C,D)** Distinct populations of *Ins2*(GFP)^LOW^ β-cells and *Ins2*(GFP)^HIGH^ β-cells were FACS purified and examined by qPCR for *gfp, Ins2,* and *Ins1* mRNA (n=3 mice). Units are relative expression calculated from ΔCT, then normalized to wild-type. One-way analysis of variance. **(E)** Dynamic imaging, over 30 hours in culture, of dispersed islet cells from *Ins1*-mCherry;*Ins2*^WT/GFP^ mice labelled with Hoechst vital nuclear dye. Note the examples of cells with bursting GFP fluorescence. Scale bar is 50 μm. * p < 0.05, ** p < 0.01, *** p < 0.001

A high-throughput, live cell imaging system with environmental control was used to study dispersed islet cells from *Ins2*^WT/GFP^:*Ins1*-mCherry mice. Remarkably, live cell imaging over ~3 days identified a sub-set of β-cells that transitioned in and out of *Ins2*(GFP)^HIGH^ activity states over the course of 36-hour long recordings (Fig. 3E, Supplemental Fig. S2A, B). Analysis of 547 cells with various cellular behavior metrics as variables identified 3 distinct clusters of *Ins2* gene activity (Supplemental Fig. S2B, C). While testing for viability of the *Ins2*^WT/GFP^ β-cells, we noticed GFP fluorescence in *Ins2*^WT/GFP^ cells sometimes rapidly decreases just prior to cell death. Therefore, in a larger study, we used propidium iodide, a dye which permits the real-time detection of the terminal stage of cell death, to identify changes in GFP fluorescence that are not associated with cell death (Fig. 4A, B, C). As propidium iodide emits red fluorescence at 636 nm, we used *Ins2*^WT/GFP^ mice without *Ins1*-mCherry for these experiments. We chose the heterozygous *Ins2*^WT/GFP^ genotype to keep the experiment as close as possible to wild-type insulin levels. Dispersed islets from *Ins2*^WT/GFP^ mice were studied for up to 4 days, imaging cells at 5-minute intervals. Similar to our pilot study, we identified β-cells that transitioned between high and low fluorescence states over the course of recordings (Fig. 4A, B). We tracked and quantified fluorescence intensities of Hoechst, GFP, propidium iodide in 2115 individual cells, confirming a bimodal distribution of mean GFP fluorescence in the cellular population (Fig. 4D, Supplemental Fig. 3A) we found in our FACS analysis. Quantification of other GFP wavelength parameters, including autocorrelation function (ACF) and full width at half maximum (FWHM) revealed groups of cells with fluctuating *Ins2* gene activity (Fig. 4E).

**Figure 4.**
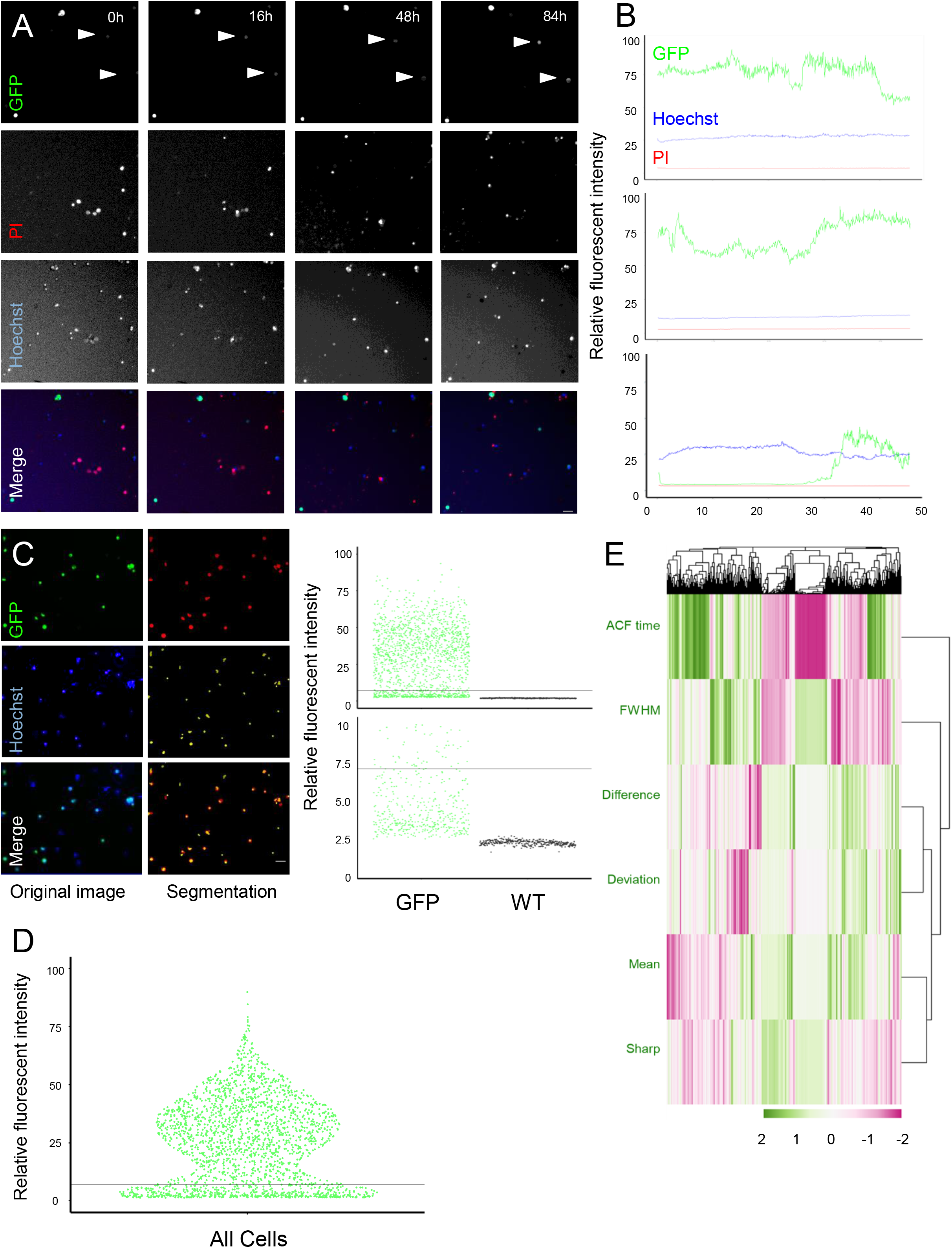
Live cell imaging of *Ins2* gene activity reveals bursting behavior of the endogenous *Ins2* gene locus. **(A)** Dynamic imaging, of up to 96 hours in culture, of dispersed islet cells from *Ins2*^WT/GFP^ mice. Cells were labelled with Hoechst vital nuclear dye and propidium iodide cell death marker. Note the examples of cells with fluctuating GFP fluorescence, indicated by white arrows. Changes in cell distribution in the red and blue wavelength can be attributed to deceased cells detaching from the bottom of the well. Scale bar is 40 μm. **(B)** Examples of cells with dynamics in relative fluorescent intensity over time. **(C)** Segmentation of cells for analysis. Cells with over 100 relative fluorescent intensity in the green wavelength compared to background were selected, comprising of ~70% of all cells imaged. This method of segmentation yields a clear cut off of cells expressing GFP compared to WT cells. Line indicates the split between high and low GFP fluorescence states. Scale bar is 40 μm. **(D)** Lifetime average GFP fluorescence of all cells imaged. Line indicates the split between high and low GFP fluorescence states. Units are relative fluorescent intensity to background. **(E)** Hierarchical clustering of cellular dynamics of *Ins2* gene activity using 6 parameters calculated from intensity over time of GFP. Parameters include deviation (mean average deviation), FWHM (full width at half maximum), mean (lifetime average GFP intensity), ACF time (function which calculates when the autocorrelation function equals 0 in a time series), difference (maximum value minus minimum value) and sharp (counts number of sharp peaks with increases greater than 10% of the mean in a cell’s lifetime). The above parameters were calculated using custom R functions. Relative fluorescent intensity: intensity relative to background, scaled 0-100.

We combined K-means clustering with principal component analysis (PCA), to reveal 4 clusters displaying distinct cellular behaviors (Supplemental Fig. S3B, C). Collectively, our long-term live cell imaging demonstrates significant dynamic fluctuations in the activity of the endogenous *Ins2* locus in primary β-cells, demonstrating that the apparently heterogeneity observed *in vivo* is not necessarily stable.

### Autocorrelation analysis of fluctuations in Ins2 gene dynamics

The observation of fluctuations in *Ins2* gene activity in some β-cells within our 4-day time window prompted us to determine the most common oscillation frequencies in *Ins2^WT/GFP^* β-cells. We performed autocorrelation analysis on GFP fluorescent intensity over time in *Ins2^WT/GFP^* β-cells that had at least one oscillation during the imaging period (Fig. 5A) and found that 357 (17%) out of 2115 cells had at least one oscillation, of which 226 had oscillation periods of 8-20 hours. The most common frequency fell within the 12-16 hour bin, with an average of ~17 hours across all oscillating cells (Fig. 5B), suggesting this oscillatory behavior may be diurnal in nature. Interestingly, we also found that cells which oscillated had significantly higher GFP fluorescent intensity in comparison to non-oscillating cells (Fig. 5B). Together, our long-term live cell imaging data suggest that circadian-influenced oscillations of *Ins2* gene activity persist in culture.

**Figure 5.**
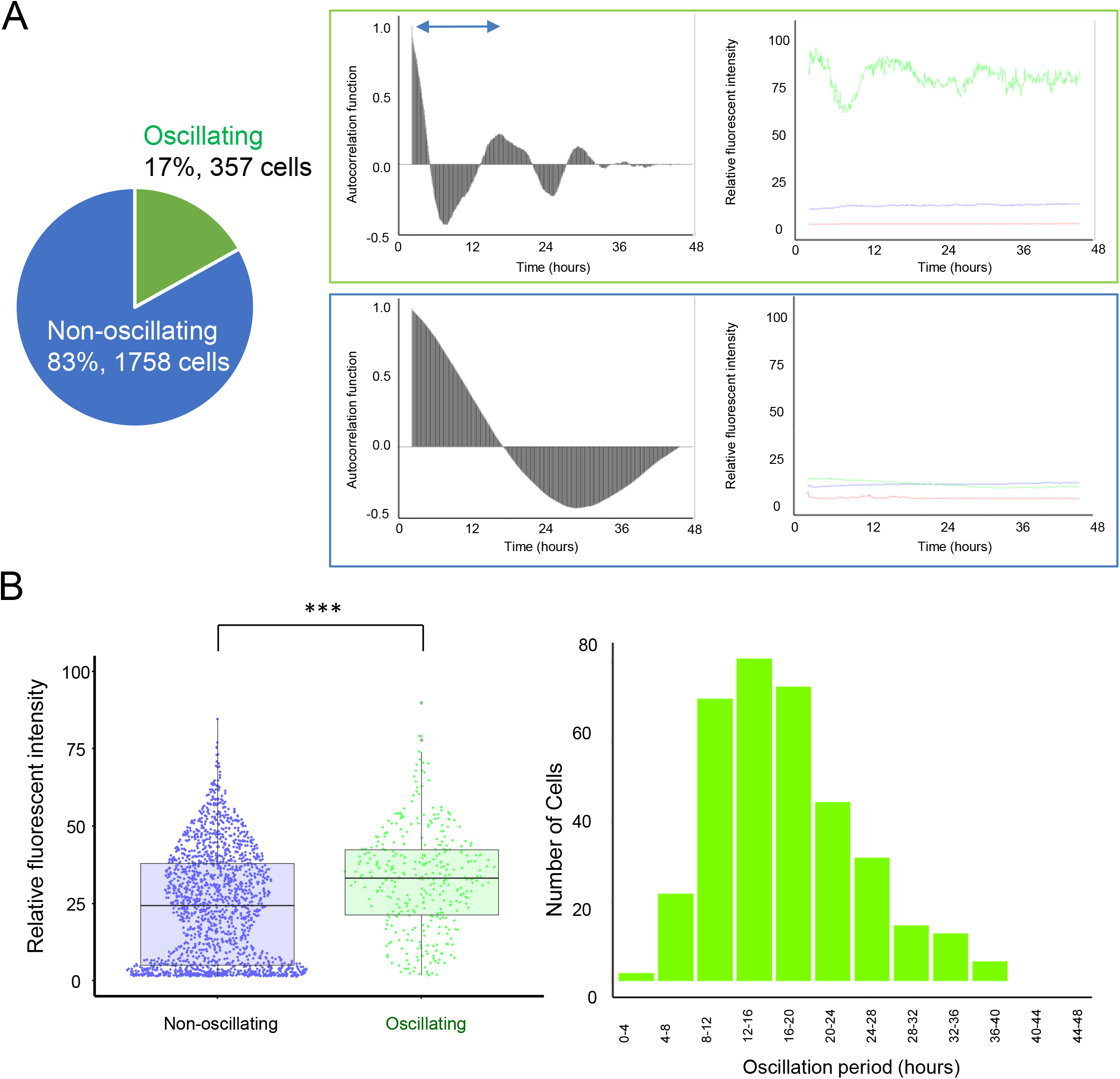
Autocorrelation analysis of cells with fluctuating behavior reveals possible connections to circadian rhythm. **(A)** Ratio of cells with at least one and no oscillations. Autocorrelation functions of cells across all time-points were calculated, and oscillation detection was done using a findpeaks function in R. For line plots, units are relative fluorescent intensity to background. For autocorrelagrams, units are autocorrelation function, calculated with the acf function in R. **(B)** Traits of oscillating cells. Average life-time GFP fluorescence intensity relative to background of oscillating versus non-oscillating cells and bar-chart of cell oscillation periods binned in 4-hour intervals. For the bar-chart, only cells with at least one oscillation were plotted (357 out of 2115 cells), and oscillation periods were calculated by finding the distance between two peaks in autocorrelograms. Relative fluorescent intensity: intensity relative to background, scaled 0-100. Student’s T-test. * p < 0.05, ** p < 0.01, *** p < 0.001

### Ins2 gene dynamics under various glucose and serum conditions

Glucose is a primary driver of insulin production, including insulin gene transcription (3; 30). Therefore, we asked if altering glucose concentrations would impact endogenous *Ins2* gene activity in our knock-in model. We treated isolated islet cells from *Ins2^WT/GFP^* mice to multiple culture conditions, including without serum (serum-free), with serum, starvation from serum for 9 hours prior to imaging (serum-starved), and an array of glucose concentrations (0, 5, 10, 15, 20 mM). The time required for mature *Ins2* mRNA production in response to elevated glucose is still under debate, with various studies describing an increase in mature *Ins2* mRNA being detected at a range of several hours after high glucose treatments, up to ~48 hours (31; 32). We found significantly higher *Ins2* gene activity (inferred by GFP fluorescence) in higher glucose concentrations under serum-containing conditions at the 48-hour mark (Fig. 6A, B). We did not find significantly higher *Ins2* gene activity in serum starved conditions, but there was still a trend towards higher gene activity in higher glucose concentrations compared to 5 mM and 10 mM glucose. In serum-free and all 0 mM glucose conditions, cells died rapidly thus confounding analysis of *Ins2* gene activity (data not shown). After de-trending our data as described in the methods, a rise in *Ins2* gene activity was observed around 12 hours into the experiment in cells cultured in higher glucose concentrations in both serum and serum-starved conditions (Fig. 6A, Supplemental Fig 4A). Interestingly, we also found a slight increase in the ratio of *Ins2*(GFP)^HIGH^ cells compared to *Ins2*(GFP)^LOW^ cells in higher glucose concentrations, which may help explain the increase in overall *Ins2* gene activity in those conditions (Supplemental Fig. S5J,). We performed hierarchical clustering with heatmap construction and extracted individual variables and compared the treatments (Supplemental Fig. S5A-I). We found a significant decrease in higher versus lower glucose concentrations in the ACF time variable (Supplemental Fig. S5H). PCA with K-means clustering analyses did not reveal any distinct clusters (Supplemental Fig. S6A, B).

**Figure 6.**
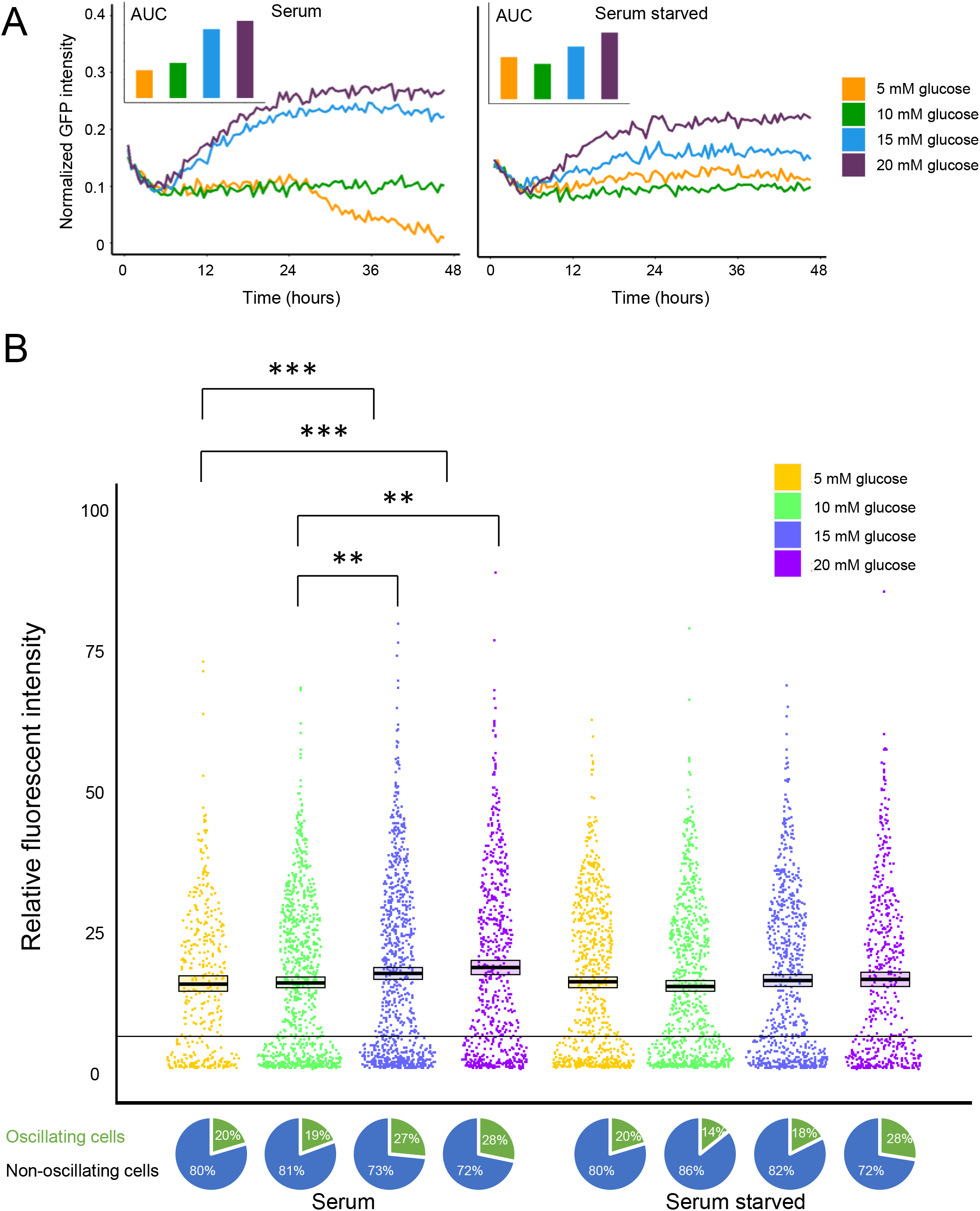
Effects of media glucose concentration on *Ins2* gene activity. **(A)** Changes in GFP fluorescence over time. Calculated by normalizing to first two hours of imaging to remove initial variation, followed by de-trending to control for photobleaching. AUC was calculated using this modified data. **(B)** Quantification and comparison of raw GFP fluorescence and ratios of oscillating versus non-oscillating cells at the 48-hour mark across all treatments. Line indicates the split between GFP high and low fluorescence states. Units are relative fluorescent intensity to background. Relative fluorescent intensity: intensity relative to background, scaled 0-100. Kruskal–Wallis one-way analysis of variance. * p < 0.05, ** p < 0.01, *** p < 0.001 WT/GFP

Autocorrelation analyses showed considerable difference in populations of oscillating versus non-oscillating cells in the various treatments. Of particular interest was the higher ratio of oscillating cells in higher glucose concentrations, suggesting that glucose may also influence *Ins2* gene expression volatility (Fig. 5B). Combined with data in our initial *Ins2^WT/GFP^* imaging study in basal 10 mM glucose conditions, this suggests that oscillating cells may be a contributor to the increase in *Ins2* gene activity in response to high glucose (Fig. 3, 4). Oscillation periods between treatments were not significantly different, but there was a trend towards higher oscillation periods in higher glucose concentrations (Supplemental Fig. S5C). We also looked at the amplitude of the oscillations by comparing the range of GFP fluorescence in oscillating cells, but we did not find any differences (Supplemental Fig. S5B). Collectively, these results show that various glucose concentrations under serum conditions may influence *Ins2* gene activity and behavior.

### Profiling β-cell states with single cell RNA sequencing

To characterize the *Ins2*(GFP)^HIGH^ state in a comprehensive and unbiased way, we performed single cell RNA sequencing on FACS purified *Ins2*(GFP)^HIGH^ and *Ins2*(GFP)^LOW^ cells from islets pooled from multiple mice (Fig. 7A-E, Supplemental Fig. S10A-D). We focused on β-cells by including only those cells expressing *Ins1* for downstream analysis (Fig 7A, B Supplemental Fig. 9A). Examination of the distribution of *Ins2* gene expression in these *Ins1*-expressing cells further confirmed our previous qPCR data (Fig. 3A) that *gfp* can accurately represent *Ins2* expression levels (Fig. 7C, Supplemental Fig. 9B). We then examined differential gene expression as a function of *gfp* mRNA (Fig. 7D, E, Supplemental Fig. S9C, D). We also considered *Ins2*(GFP)^HIGH^ and *Ins2*(GFP)^LOW^ as binary categories and examined differential gene expression (Supplemental Fig. S8 A, B).

**Figure 7.**
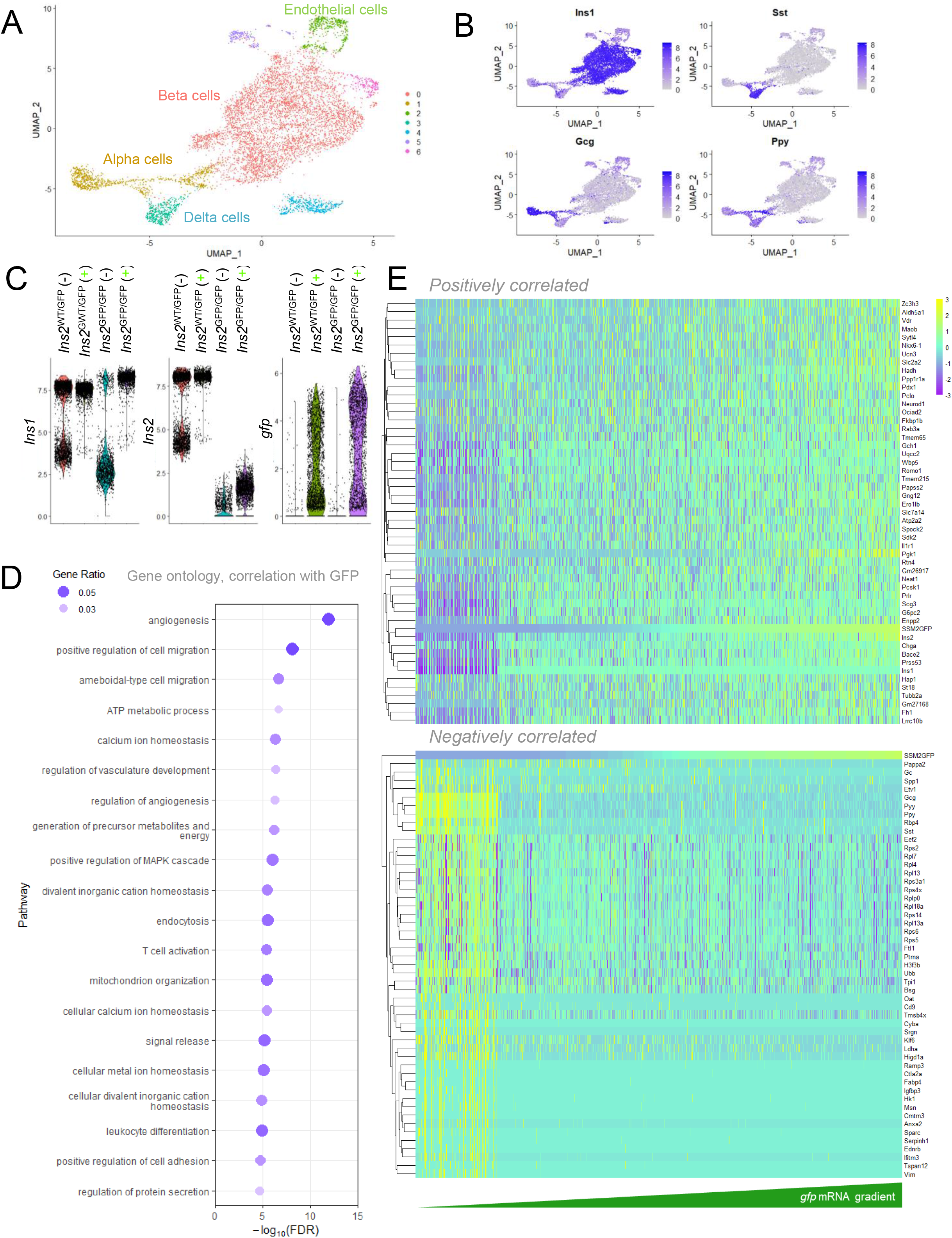
Single cell RNA sequencing analysis of *Ins2*(GFP)^LOW^ and *Ins2*(GFP)^HIGH^ β-cells in 60-week-old mice. **(A)** UMAP plot of all cells isolated and dispersed from *Ins2*^WT/GFP^ or *Ins2*^GFP/GFP^ mice and FACS purified into either GFP-negative (-) or GFP-positive (+) groups. **(B)** UMAPs projections with known markers for major cell populations in islets. Cells were selected for analysis based on high *Ins1* expression and low expression of other cell population markers. **(C)** Log-normalized counts of *Ins1, Ins2,* and *gfp* mRNA quantification distributions from GFP-negative or GFP-positive single β-cells. **(D)** Gene Ontology categories that are driven by genes differentially expressed in *Ins2*(GFP)^LOW^ versus *Ins2*(GFP)^HIGH^ β-cells from old heterozygous *Ins2*^WT/GFP^ mice. **(E)** Heatmap showing **i** ndividual genes that are differentially expressed as a function of *gfp* mRNA expression in old heterozygous *Ins2*^WT/GFP^ mice.

In 8-week-old *Ins2*^WT/GFP^ mice, pathway enrichment analysis of cells expressing high *gfp* showed that the *Ins2*(GFP)^HIGH^ cells had significant alterations in genes involved in hormone regulation, hormone secretion, hormone transport, protein synthesis, and protein cleavage (Supplemental Fig. S9C). Cluster of genes ordered by correlation with *gfp* in heterozygous *Ins2*^WT/GFP^ β-cells showed that *Sema6a,* and the antioxidant metallothionein genes *Mt1* and *Mt2,* were most closely correlated to *gfp* mRNA at the single cell level. Classical genes related to β-cell function and maturity that were positively correlated with *gfp* include *Ins2, Ero1lb,* Chromogranin A (*Chga*), the Glut2 glucose transporter *Slc2a2, Iapp, Nkx6-1,* and *Pdx1* (18; 33; 34) (Supplemental Fig. 9D). The *gfp-*high state was also characterized by increased mRNA expression of genes encoding signal recognition particle 9 (*Srp9*), G-protein subunit gamma 12 (*Gng12*), nucleotide pyrophosphatase/phosphodiesterase 2 (*Enpp2*), aldehyde dehydrogenase 5 family member A1 (*Aldh5a1*), and tetraspanin-28 (*Cd81).* Stress response genes, such as *Sec61b, Sec61g* and *Sec23b* were also highly expressed (35; 36) (Supplemental Fig. 9D). The *gfp-*high state was associated with decreased expression of genes including soluble factors such as peptide YY (*Pyy*), somatostatin (*Sst*) glucagon (*Gcg*) pancreatic polypeptide (*Ppy*), suggesting in aggregate a less ‘poly-hormonal’ and therefore, more mature, gene expression profile (17; 37) (Supplemental Fig. 9D). In agreement, *gfp-*high β-cells had lower expression of pappalysin 2 (*Pappa2*), an α cell-selective regulator of insulin-like growth factor bioavailability, and the epsilon cell marker (*Etv1)* (38) (Supplemental Fig. S9D). *Gfp-*high cells also had reduced expression of several genes linked to insulin production and secretion, such as multiple subunits of the 40S and 60S ribosomes, eukaryotic translation initiation factor 3 subunit C (*Eif3c*), Eukaryotic translation initiation factor 2 subunit 2 (*Eif2s2*), and heat shock 70kDa protein 9 (*Hspa9;* Mortalin) (Supplemental Fig. S9D). Interestingly, a decrease in the expression of certain ER stress genes was found, including *Prkcsh, Nucb2,* and *Fam129a* (39–41) (Supplemental Fig. S9D). Thus, in young mice, the *Ins2*(GFP)^HIGH^ cell state is associated with a mature single cell gene expression profile, optimal insulin secretion, and a reorganization of protein synthesis machinery.

Age is known to significantly alter the properties of pancreatic β-cells, including their function and ability to enter the cell cycle (42). Thus, we conducted an additional similar study in three 60-week-old mice. In this experiment, we studied cells from both *Ins2*^WT/GFP^ (Fig. 7A-E) and *Ins2*^GFP/GFP^ islets (Supplemental Fig. S10A-C). Pathway enrichment analysis of cells expressing high *gfp* showed that the *Ins2*(GFP)^HIGH^ cells had significant alterations in genes involved in vasculature development, metabolism, and ion homeostasis (Fig. 7D). Cluster analysis of genes ordered by *gfp* gradient in old heterozygous *Ins2*^WT/GFP^ β-cells revealed that *Ins2* expression most closely matches *gfp* mRNA (Fig. 7E). Similar to the younger mice, genes that increased with *gfp* expression included genes related to optimal insulin secretion such as *Chga* and *Slc2a2,* as well as key β-cell transcription factors and maturity markers *Nkx6-1* and *Pdx1* (Fig. 7E). Other genes related to insulin production and secretion that were upregulated include *Ins1,* secretogranin 3 (*Scg3*), proprotein convertase subtilisin/kexin type 1 (*Pcsk1*), ubiquinol-cytochrome c reductase complex assembly factor 2 (*Uqcc2*), synaptotagmin-like 4 (*Sytl4)* (25; 43; 44) (Fig. 7E). There was also an enrichment in key β-cell transcription factors and maturity markers *Ucn3* and *Neurod1* (45; 46) (Fig. 7E). Interestingly, many of the mRNAs that were increased are known to be *Pdx1* target genes in islets (47). Other notable genes that were upregulated in *gfp*-high cells were *Gng12, Enpp2, Ppp1r1a,* and metabolism-regulating genes, such as glucose-6-phosphatase catalytic subunit 2 (*G6pc2)* and hydroxyacyl-CoA dehydrogenase (*Hadh)* (Fig. 7E). ER stress-related gene *Neat1* was also increased (48). Also in agreement with the analysis of young islets, *gfp*-high β-cells from old homozygous mice had decreased expression of *Pappa2, Etv1,* and *Sst*, as well as other markers of β-cell immaturity (*Gcg, Ppy, Pyy). Ldha,* another dedifferentiation marker, was also negatively correlated with *gfp* (Fig. 7E). Several ER stress genes were also anti-correlated with *gfp,* including *Eef2* and *Spp1* (49; 50) (Fig. 7E). Thus, in older mice, the *Ins2*(GFP)^HIGH^ cell state is overall similar to younger mice, with a more mature single cell expression profile and association with genes related to metabolism.

Separately, we analyzed genes ordered by *gfp* mRNA gradient in old homozygous *Ins2*^GFP/GFP^ β-cells (Supplemental Fig. S10). Many of the same genes that we observed in the heterozygous samples had similar expression patterns in the GFP homozygous samples, including maturity markers *Chga, Neurod1, Pdx1, Ucn3, Slc2a2, Gng12,* and *Ppp1r1a,* and de-differentiation markers *Sst*, *Pyy, Ppy, Gcg.* Nuclear protein 1, transcriptional regulator (*Nupr1*), a stress adaptation gene, was reduced in homozygous *Ins2*^GFP/GFP^ β-cells with high *gfp* expression. When we combined data from old *Ins2*^WT/GFP^ and *Ins2*^GFP/GFP^ cells, pathway enrichment analysis of high *gfp*-expressing cells in older mice showed that the *Ins2*(GFP)^HIGH^ cells had significant alterations in genes involved in protein translation, RNA splicing, mRNA processing (Supplemental Fig. S10B). With this combined analysis, *gfp* correlation with many of the same genes found by analysing *Ins2*^WT/GFP^ and *Ins2*^GFP/GFP^ cells separately were also identified (Supplemental Fig. S11). Pathway enrichment analysis of differential expression in all *Ins1* positive cells revealed upregulation in genes related to translation machinery in younger mice, while in older mice there was upregulation in genes related to cellular respiration (Supplemental Fig. S12A, B).

The single cell RNA sequencing studies identified enrichment in genes that control β-cell stress responses and markers of β-cell maturity. We followed this up by performing RNA velocity analysis on older *Ins2*^WT/GFP^ and *Ins2*^GFP/GFP^ mice. We found that the velocity vectors indicate β-cells of older *Ins2*^WT/GFP^ and *Ins2*^GFP/GFP^ with higher levels of *gfp* mRNA were more mature than β-cells with lower levels of *gfp* mRNA (Fig 8A, Supplemental Fig. S12A). Due to high dropout rates of *gfp* mRNA in the younger mice, we were unable to perform RNA velocity analysis on those mice.

**Figure 8.**
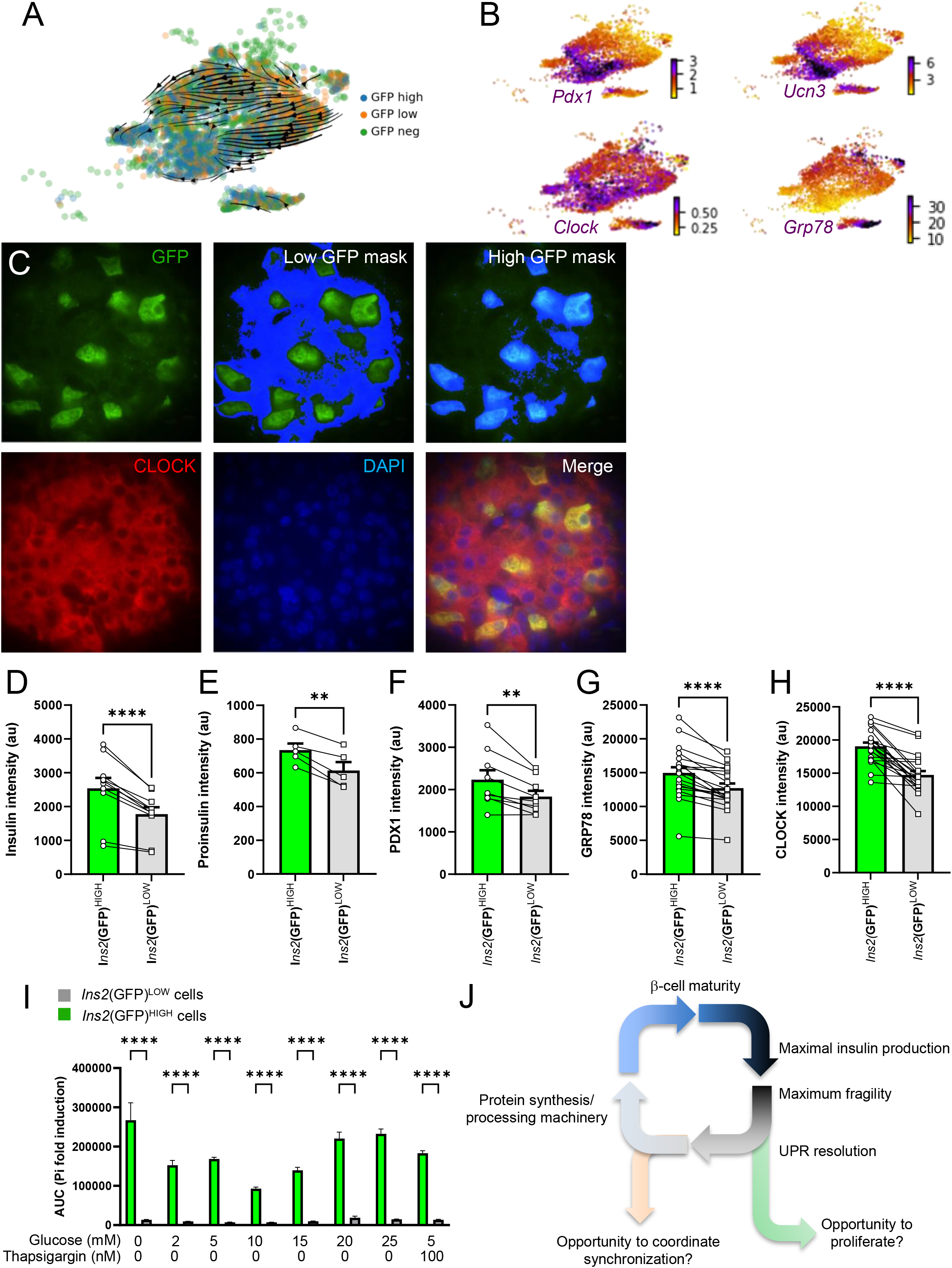
*Ins2*(GFP)^HIGH^ β-cells show increased markers of maturity and increased cell death across multiple conditions. **(A)** UMAP projection of β-cells with RNA velocity vectors overlaid. **(B)** Expression levels of genes of interest. Genes selected include genes which serve as indicators of β-cell maturity, the circadian cycle, and ER stress. **(C)** Immunofluorescent staining of pancreatic sections from *Ins2*^WT/GFP^ mice reveals increased expression in markers of maturity, circadian cycle, and ER stress in high GFP expressing cells. Proteins stained were chosen based on expression in single cell RNA sequencing data. Quantification was done by arbitrarily selecting for regions with high GFP fluorescence and comparing to regions with low GFP fluorescence. **(D-H)** Quantification of immunofluorescent staining for insulin, proinsulin, PDX1, GRP78, and CLOCK. **(I)** Cells with high GFP intensities show increased apoptosis susceptibility at all tested glucose concentrations and in the presence of thapsigargin (>1000 cells/condition, repeated from 2 mice). **(J)** Hypothetical working model of transitions between insulin production states in β-cells.

Our single cell analyses also showed correlation of *gfp* with various factors related to optimal insulin secretion and processing. Immunofluorescent staining on insulin and proinsulin indicated increased expression of both in high GFP-expressing cells, hinting at a possible difference in functionality between the β-cell states (Fig. 8 D, E). We further investigated the maturation aspect by performing immunofluorescent staining on select maturation markers which were upregulated in high *gfp*-expressing cells. We found that PDX1 levels were significantly increased in high GFP-expressing cells compared with lower GFP-expressing cells (Fig. 8F). However, we did not find a significant difference in CHGA, another maturity marker (Supplemental Fig. 14C). We next investigated individual gene products in an attempt to elucidate pathways that are upregulated in high GFP-expressing cells (Supplemental Fig. 11). We have previously shown that insulin production itself is a significant stress under basal conditions in β-cells (1) and we therefore predicted that cells with increased *Ins2* gene activity and GFP production would be more sensitive to stress. Interestingly, we found that *Hspa5* (GRP78), a major ER stress regulator, was downregulated in high *gfp*-expressing cells (Fig. 8G). Unexpectedly, immunofluorescent staining on pancreatic sections obtained from *Ins2*^WT/GFP^ mice showed that GRP78 protein levels were upregulated in high GFP-expressing cells. The autocorrelation analyses revealed possible connections to circadian cycles. Indeed, high *gfp*-expressing cells had increased levels of *Clock* gene expression compared to lower *gfp*-expressing cells (Fig. 8B). Immunofluorescent staining for CLOCK protein showed that high GFP-expressing cells had higher levels of CLOCK protein than lower GFP-expressing cells, confirming our single cell analyses (Fig. 8H). Thus, the states marked by high gene activity at the endogenous *Ins2* locus likely possess critical functional differences.

Finally, we asked if high insulin production was associated with increased β-cell fragility. Indeed, using live cell imaging, we found that *Ins2*(GFP)^HIGH^ cells were >10 fold more sensitive to apoptosis at all glucose concentrations we tested when compared with *Ins2*(GFP)^LOW^ cells in the same cultures (Fig. 8I). Collectively, we have shown that insulin gene activity marks β-cell states, with consequences for function and survival.

## Discussion

The goal of the present study was to determine the nature of *Ins2* gene expression heterogeneity. Analysis of pancreatic tissue sections from *Ins2*^GFP^ knock-in mice showed that, at any given time, roughly 40% of all β-cells exist in a GFP-high state, suggesting that not all β-cells have simultaneously high levels of active transcription at the *Ins2* locus *in vivo.* Over the course of multi-day *in vitro* imaging experiments, we observed clear transitions between *Ins2*(GFP)^HIGH^ and *Ins2*(GFP)^LOW^ states in single β-cells, possibly linked to the circadian clock. However, *Ins2* gene activity was stable for the duration of these studies in the majority of cells. We used single cell RNA sequencing to characterize the *Ins2*(GFP)^HIGH^ cellular state and found that *Ins2*(GFP)^HIGH^ were significantly more sensitive under all stress conditions examined. Together with previous live cell imaging data, the results of the present study demonstrate that a substantial component of β-cell heterogeneity in relation to insulin gene expression is dynamic in time (Fig. 8J).

The dynamics of GFP fluorescence revealed by live cell imaging of dispersed islet cells from *Ins2*^GFP^ mice provided an unprecedented look at insulin gene activity in populations of single β-cells. Many of the *Ins2*(GFP)^LOW^ and *Ins2*(GFP)^HIGH^ β-cell states were maintained over at least 24 hours. Autocorrelation analysis showed that of the cells which displayed at least one oscillation, a significant proportion have oscillation periods clustered at 8-12 hours. There were significantly more GFP^HIGH^ cells with oscillations compared with GFP^LOW^ cells. We also found a significant increase of Clock gene and protein levels in high GFP-expressing cells. Circadian control of islet function is essential for maintaining normal glucose homeostasis, and circadian-based variations on the transcriptional level have been described in diverse cell types and shown to be critical for optimization of cellular function (51). Circadian influences are known to persist in cell culture in β-cells, and insulin is both a target of circadian regulation, and a regulator of the circadian machinery in many cell types (52–54). Diurnal rhythms of β-cell function and metabolism has long been recognized (55; 56), and our study extends this to activity at the insulin gene locus.

The study of pancreatic islet cell heterogeneity is currently experiencing resurgence, in part due to the application of single cell sequencing and optogenetic technologies to answer islet biology questions. There are many examples of β-cell heterogeneity and these were reviewed recently (4). Insulin gene expression is a cardinal feature of pancreatic β-cells, but cell-by cell variability in insulin production has remained under-appreciated despite published evidence (11). For example, there are reports of significant β-cell heterogeneity in transgenic mice expressing GFP under the *Ins2* promoter (57). Similarly, we have shown variation in fluorescent protein expression under the control of *Ins1* promoters *in vivo* and *in vitro* (13–15). However, a limitation in these studies is that artificial promoter constructs may not recapitulate the more complex and long-range regulation available at the endogenous gene locus. Notwithstanding, cell-by cell analysis of insulin mRNA, either by singlemolecule fluorescent *in situ* hybridization or single cell RNA sequencing, also showed a 2-to 10-fold range in native gene expression from endogenous insulin gene loci (8; 12). The *Ins2*(GFP)^HIGH^ β-cells we identified in our study are possibly a temporal manifestation of the extreme β-cells reported by Farack et al (12).For instance, their extreme β-cells had significantly elevated *Chga, Pdx1, Slc2a2,* and *Ucn3* mRNA expression (12), which is consistent with our single cell RNA sequencing data. It is possible that the dynamic nature of insulin transcription is an adaptation to the extremely high demands of producing and maintaining adequate stores of insulin in β-cells. Assuming there are homeostatic mechanisms to maintain stable insulin protein stores, it is unclear how the need for a burst of insulin gene activity would be sensed by individual β-cells, and future studies should attempt to define these mechanisms.

The relationship between β-cell states demarked by insulin expression and disease pathogenesis remains unclear. Type 1 diabetes risk is associated with an increase in *INS* mRNA in whole human fetal pancreas (58) and single adult β-cells pancreas (59). Isolated islets and single human β-cells from people with type 2 diabetes were reported to have reduced *INS* expression on average (38; 60), although our recent meta-analysis of single cell RNA sequencing data sets did not find a statistically significant reduction in INS mRNA levels in β-cells from donors with type 2 diabetes (59). Extreme β-cells were significantly more common in diabetic *db/db* mice (12). We found that the *Ins2*(GFP)^HIGH^ state was associated with significant vulnerability to cellular stress, so having an excess number of *Ins2*(GFP)^HIGH^ β-cells at a given time may negatively affect islet health and robustness. These results are consistent with our previous data defining the interrelationships between maximal insulin production, ER stress and β-cell proliferation (1). We also note that in more hyperglycemic conditions, more cells may start to transfer towards the *Ins2*(GFP)^HIGH^ state, but due to the small timeframe that the imaging allows, we were unable to consistently observe cells transition from *Ins2*(GFP)^LOW^ to the higher end of *Ins2*(GFP)^HIGH^ states. It has been shown that rest may be important for β-cells, as dysfunctional cells that were removed from diabetic mice were shown to recover functionality when cultured at normal glucose levels (3). Thus, the ability to transition back to *Ins2*(GFP)^LOW^ states may also be critical, as it may represent a form of rest for the β-cells, with failure in doing so possibly resulting in chronic stress and dysfunction. It is also likely that stress may modulate the frequency of β-cell state transitions, although we did not test this directly.

The presence of a proportion of β-cells in the *Ins2*(GFP)^LOW^ state may be essential for islet function, with recent studies showing that loss of less mature β-cells impacts overall islet function (61). Specifically, so called ‘hub β-cells’ that help synchronize islets were reported to have lower insulin content compared with typical β-cells (5), and we have speculated that this represents a trade-off needed for their synchronizing function (6). Rodent and human β-cells are long-lived (62; 63), and perhaps β-cells cycle through multiple states during their existence, including taking turns supporting the oscillatory coupling of the islet. Given that robust β-cell heterogeneity and state transitions have not been reported for stem cell derived β-like cells (64), it is likely that *in vitro* differentiation protocols will need to be further optimized to produce a full range of dynamic β-cell characteristics. Interestingly, many of the genes that are differentially expressed in *Ins2*(GFP)^HIGH^ β-cells are known to play roles in type 2 diabetes susceptibility, including common alleles of the MODY/neonatal diabetes genes *Pdx1, Neurod1, Nkx6.1, Abcc8, Slc2a2,* as well as *Slc30a8* and *Pam* previously identified by genome-wide association (65). It will also be interesting to examine the frequency of β-cell states in the context of type 1 diabetes, given that pro-insulin, *Pdx1, Iapp,* and *Chga* are auto-antigens (66–68). Indeed, β-cells undergoing proliferation or with lower insulin/maturity are protected in the NOD mouse model of type 1 diabetes (69; 70). Collectively, these observations suggest that modulation of β-cell states could be a therapeutic and/or prevention target for both type 1 diabetes and type 2 diabetes.

Temporal transcriptional plasticity and gene expression bursting on a similar time scale as to what we have observed have been documented in bacteria, yeast and other mammalian cell types (71–73). For example, bursting gene expression patterns have been observed in pituitary cells (74–76). Interestingly, LHβ transcription in pituitary gonadotrophs is directly linked to proteasome activity (75), suggesting a possible mechanism for coupling protein loads and transcription in secretory cell types. Many cell-extrinsic and cell-intrinsic factors have been implicated in the modulation of transcriptional burst frequency, including histone modifications and chromatin topology (77–81). Future studies will be required to determine the molecular mechanisms mediating transcriptional bursting at the insulin gene locus in β-cells. Future studies should also seek to directly measure *Ins2* mRNA transcription, perhaps using new CRISPR-based probes or other mRNA tagging systems (82–84).

In this study, we identified a subpopulation of cells that had relatively rapid transitions between *Ins2*(GFP)^LOW^ and *Ins2*(GFP)^HIGH^ β-cell states suggesting dynamic transcriptional activity at the *Ins2* gene locus rather than stable heterogeneity. However, our study has several limitations. For example, our measurements of GFP fluorescence originating from the *Ins2* locus*-*mediated transcription cannot distinguish the relative contribution in changes in GFP mRNA transcription/stability or GFP protein translation/degradation. The half-life of unmodified GFP is ~26 hours (85) and the changes in GFP fluorescence in some cells suggest coordinated increases of protein synthesis and coupled protein degradation, which may also explain the slow decrease observed in our over time analyses. Also, oscillations detected in the *in vitro* live cell imaging experiments cannot be used to determine the ratio of oscillating cells *in vivo,* due to the possible loss of intercellular signals and additional stress in imaging dispersed cells. While some intercellular signals could be missing in our analyses of dispersed islet cells (86; 87), our *in vivo* analysis revealed a similar proportion of *Ins2*(GFP)^LOW^ and *Ins2*(GFP)^HIGH^ giving us confidence that the two-population phenomenon we measured *in vitro* is generalizable to *in vivo* conditions. Future experiments should involve more powerful live cell imaging techniques, including tissue slides and intravitreal imaging, ideally with techniques that allow long-term cell tracking without excess phototoxicity or photobleaching (88; 89).

In conclusion, our data demonstrate that single β-cells can switch between states marked by high and low activity of the phylogenetically conserved, endogenous insulin gene locus. This newly discovered phenomenon may account for much of the observed heterogeneity in β-cell insulin gene expression measured at single time points and needs to be comprehensively studied and leveraged in efforts to protect β-cells in the context of diabetes and to generate new β-cells from stem cells (17; 90).

## Author Contributions

CMJC designed studies, performed experiments, analyzed/interpreted data, wrote the revised manuscript. HM designed studies, performed experiments, analyzed/interpreted data, and wrote the original version of the manuscript. SS performed experiments, analyzed/interpreted data. CE analyzed/interpreted data. HC performed experiments, analyzed/interpreted data. NAJK analyzed/interpreted data. DAD performed experiments, analyzed/interpreted data. BX analyzed/interpreted data. XH performed experiments. NN performed experiments and analyzed/interpreted data. EP supervised work and analyzed/interpreted data. YX analyzed/interpreted data. SX provided a key reagent. MOH provided a key reagent, designed studies, and analyzed/interpreted data. TJK supervised work. FCL supervised work, analyzed/interpreted data, and edited the manuscript. JDJ conceived the project, designed studies, interpreted data, edited the manuscript, and is the guarantor of this work.

## Declaration of Interests

The authors declare no competing interests.

## Funding

Research was supported by a CIHR operating grant (PJT-152999) to J.D.J. and the JDRF Centre of Excellence at UBC.

## Acknowledgments

We thank many colleagues for helpful discussions. This manuscript used data acquired from the Human Pancreas Analysis Program (HPAP-RRID:SCR_016202) Database (https://hpap.pmacs.upenn.edu), a Human Islet Research Network (RRID:SCR_014393) consortium (UC4-DK-112217, U01-DK-123594, UC4-DK-112232, and U01-DK-123716).

## Notes

### Competing Interest Statement

The authors have declared no competing interest.

### Summary of Updates

New data, including additional verification on certain topics and follow-up studies on differentially expressed genes identified in scRNAseq data.

## References

1. Szabat M, Page MM, Panzhinskiy E, Skovso S, Mojibian M, Fernandez-Tajes J, Bruin JE, Bround MJ, Lee JT, Xu EE, Taghizadeh F, O’Dwyer S, van de Bunt M, Moon KM, Sinha S, Han J, Fan Y, Lynn FC, Trucco M, Borchers CH, Foster LJ, Nislow C, Kieffer TJ, Johnson JD: Reduced Insulin Production Relieves Endoplasmic Reticulum Stress and Induces beta Cell Proliferation. Cell Metab 2016;23:179–193

2. Wang S, Flibotte S, Camunas-Soler J, MacDonald PE, Johnson JD: A New Hypothesis for Type 1 Diabetes Risk: The At-Risk Allele at rs3842753 Associates With Increased Beta-cell INS Messenger RNA in a Meta-Analysis of Single-Cell RNA-Sequencing Data. Can J Diabetes 2021;

3. Boland BB, Rhodes CJ, Grimsby JS: The dynamic plasticity of insulin production in beta-cells. Mol Metab 2017;6:958–973

4. Benninger RKP, Hodson DJ: New Understanding of beta-Cell Heterogeneity and In Situ Islet Function. Diabetes 2018;67:537–547

5. Johnston NR, Mitchell RK, Haythorne E, Pessoa MP, Semplici F, Ferrer J, Piemonti L, Marchetti P, Bugliani M, Bosco D, Berishvili E, Duncanson P, Watkinson M, Broichhagen J, Trauner D, Rutter GA, Hodson DJ: Beta Cell Hubs Dictate Pancreatic Islet Responses to Glucose. Cell Metab 2016;24:389–401

6. Kolic J, Johnson JD: Specialized Hub Beta Cells Trade Maximal Insulin Production for Perfect Timing. Cell Metab 2016;24:371–373

7. Dorrell C, Schug J, Canaday PS, Russ HA, Tarlow BD, Grompe MT, Horton T, Hebrok M, Streeter PR, Kaestner KH, Grompe M: Human islets contain four distinct subtypes of beta cells. Nat Commun 2016;7:11756

8. Xin Y, Dominguez Gutierrez G, Okamoto H, Kim J, Lee AH, Adler C, Ni M, Yancopoulos GD, Murphy AJ, Gromada J: Pseudotime Ordering of Single Human beta-Cells Reveals States of Insulin Production and Unfolded Protein Response. Diabetes 2018;67:1783–1794

9. Wills QF, Boothe T, Asadi A, Ao Z, Warnock GL, Kieffer TJ, Johnson JD: Statistical approaches and software for clustering islet cell functional heterogeneity. Islets 2016;8:48–56

10. Pipeleers DG: Heterogeneity in pancreatic beta-cell population. Diabetes 1992;41:777–781

11. Kiekens R, In ‘t Veld P, Mahler T, Schuit F, Van De Winkel M, Pipeleers D: Differences in glucose recognition by individual rat pancreatic B cells are associated with intercellular differences in glucose-induced biosynthetic activity. J Clin Invest 1992;89:117–125

12. Farack L, Golan M, Egozi A, Dezorella N, Bahar Halpern K, Ben-Moshe S, Garzilli I, Toth B, Roitman L, Krizhanovsky V, Itzkovitz S: Transcriptional Heterogeneity of Beta Cells in the Intact Pancreas. Dev Cell 2019;48:115–125 e114

13. Szabat M, Luciani DS, Piret JM, Johnson JD: Maturation of adult beta-cells revealed using a Pdx1/insulin dual-reporter lentivirus. Endocrinology 2009;150:1627–1635

14. Szabat M, Pourghaderi P, Soukhatcheva G, Verchere CB, Warnock GL, Piret JM, Johnson JD: Kinetics and genomic profiling of adult human and mouse beta-cell maturation. Islets 2011;3:175–187

15. Szabat M, Johnson JD, Piret JM: Reciprocal modulation of adult beta cell maturity by activin A and follistatin. Diabetologia 2010;53:1680–1689

16. Wakae-Takada N, Xuan S, Watanabe K, Meda P, Leibel RL: Molecular basis for the regulation of islet beta cell mass in mice: the role of E-cadherin. Diabetologia 2013;56:856–866

17. Johnson JD: The quest to make fully functional human pancreatic beta-cells from embryonic stem cells: Climbing a mountain in the clouds. Diabetologia 2016;59:2047–2057

18. Benner C, van der Meulen T, Caceres E, Tigyi K, Donaldson CJ, Huising MO: The transcriptional landscape of mouse beta cells compared to human beta cells reveals notable species differences in long non-coding RNA and protein-coding gene expression. BMC Genomics 2014;15:620

19. Kaestner KH, Powers AC, Naji A, Consortium H, Atkinson MA: NIH Initiative to Improve Understanding of the Pancreas, Islet, and Autoimmunity in Type 1 Diabetes: The Human Pancreas Analysis Program (HPAP). Diabetes 2019;68:1394–1402

20. Yang YH, Johnson JD: Multi-parameter single-cell kinetic analysis reveals multiple modes of cell death in primary pancreatic beta-cells. J Cell Sci 2013;126:4286–4295

21. Matthews DR, Lang DA, Burnett MA, Turner RC: Control of pulsatile insulin secretion in man. Diabetologia 1983;24:231–237

22. Scrucca L, Fop M, Murphy TB, Raftery AE: mclust 5: Clustering, Classification and Density Estimation Using Gaussian Finite Mixture Models. R J 2016;8:289–317

23. Chen S, Thakor NV, Mower MM: Ventricular fibrillation detection by a regression test on the autocorrelation function. Med Biol Eng Comput 1987;25:241–249

24. Butler A, Hoffman P, Smibert P, Papalexi E, Satija R: Integrating single-cell transcriptomic data across different conditions, technologies, and species. Nat Biotechnol 2018;36:411–420

25. Szabat M, Lynn FC, Hoffman BG, Kieffer TJ, Allan DW, Johnson JD: Maintenance of beta-cell maturity and plasticity in the adult pancreas: developmental biology concepts in adult physiology. Diabetes 2012;61:1365–1371

26. Johnson JD, Ahmed NT, Luciani DS, Han Z, Tran H, Fujita J, Misler S, Edlund H, Polonsky KS: Increased islet apoptosis in Pdx1+/-mice. J Clin Invest 2003;111:1147–1160

27. Johnson JD, Bernal-Mizrachi E, Alejandro EU, Han Z, Kalynyak TB, Li H, Beith JL, Gross J, Warnock GL, Townsend RR, Permutt MA, Polonsky KS: Insulin protects islets from apoptosis via Pdx1 and specific changes in the human islet proteome. Proc Natl Acad Sci U S A 2006;103:19575–19580

28. Mehran AE, Templeman NM, Brigidi GS, Lim GE, Chu KY, Hu X, Botezelli JD, Asadi A, Hoffman BG, Kieffer TJ, Bamji SX, Clee SM, Johnson JD: Hyperinsulinemia drives diet-induced obesity independently of brain insulin production. Cell Metabolism 2012;16:723–737

29. Leroux L, Desbois P, Lamotte L, Duvillie B, Cordonnier N, Jackerott M, Jami J, Bucchini D, Joshi RL: Compensatory responses in mice carrying a null mutation for Ins1 or Ins2. Diabetes 2001;50 Suppl 1:S150–153

30. Knoch KP, Nath-Sain S, Petzold A, Schneider H, Beck M, Wegbrod C, Sonmez A, Munster C, Friedrich A, Roivainen M, Solimena M: PTBP1 is required for glucose-stimulated cap-independent translation of insulin granule proteins and Coxsackieviruses in beta cells. Mol Metab 2014;3:518–530

31. Evans-Molina C, Garmey JC, Ketchum R, Brayman KL, Deng S, Mirmira RG: Glucose regulation of insulin gene transcription and pre-mRNA processing in human islets. Diabetes 2007;56:827–835

32. Tillmar L, Carlsson C, Welsh N: Control of insulin mRNA stability in rat pancreatic islets. Regulatory role of a 3’-untranslated region pyrimidine-rich sequence. J Biol Chem 2002;277:1099–1106

33. Zito E, Chin KT, Blais J, Harding HP, Ron D: ERO1-beta, a pancreas-specific disulfide oxidase, promotes insulin biogenesis and glucose homeostasis. J Cell Biol 2010;188:821–832

34. Augsornworawat P, Maxwell KG, Velazco-Cruz L, Millman JR: Single-cell transcriptome profiling reveals beta cell maturation in stem cell-derived islets after transplantation. Cell Rep 2021;34:108850

35. Sundaram A, Plumb R, Appathurai S, Mariappan M: The Sec61 translocon limits IRE1alpha signaling during the unfolded protein response. Elife 2017;6

36. Yehia L, Liu D, Fu S, Iyer P, Eng C: Non-canonical role of wild-type SEC23B in the cellular stress response pathway. Cell Death Dis 2021;12:304

37. Basford CL, Prentice KJ, Hardy AB, Sarangi F, Micallef SJ, Li X, Guo Q, Elefanty AG, Stanley EG, Keller G, Allister EM, Nostro MC, Wheeler MB: The functional and molecular characterisation of human embryonic stem cell-derived insulin-positive cells compared with adult pancreatic beta cells. Diabetologia 2012;55:358–371

38. Segerstolpe A, Palasantza A, Eliasson P, Andersson EM, Andreasson AC, Sun X, Picelli S, Sabirsh A, Clausen M, Bjursell MK, Smith DM, Kasper M, Ammala C, Sandberg R: Single-Cell Transcriptome Profiling of Human Pancreatic Islets in Health and Type 2 Diabetes. Cell Metab 2016;24:593–607

39. Shin GC, Moon SU, Kang HS, Choi HS, Han HD, Kim KH: PRKCSH contributes to tumorigenesis by selective boosting of IRE1 signaling pathway. Nat Commun 2019;10:3185

40. Zhang D, Lin J, Chao Y, Zhang L, Jin L, Li N, He R, Ma B, Zhao W, Han C: Regulation of the adaptation to ER stress by KLF4 facilitates melanoma cell metastasis via upregulating NUCB2 expression. J Exp Clin Cancer Res 2018;37:176

41. Evstafieva AG, Kovaleva IE, Shoshinova MS, Budanov AV, Chumakov PM: Implication of KRT16, FAM129A and HKDC1 genes as ATF4 regulated components of the integrated stress response. PLoS One 2018;13:e0191107

42. Rankin MM, Kushner JA: Adaptive beta-cell proliferation is severely restricted with advanced age. Diabetes 2009;58:1365–1372

43. Cambier L, Rassam P, Chabi B, Mezghenna K, Gross R, Eveno E, Auffray C, Wrutniak-Cabello C, Lajoix AD, Pomies P: M19 modulates skeletal muscle differentiation and insulin secretion in pancreatic beta-cells through modulation of respiratory chain activity. PLoS One 2012;7:e31815

44. Lin CC, Cheng KP, Hung HC, Li CH, Lin CH, Chang CJ, Hu CY, Wu HT, Ou HY: Serum Secretogranin III Concentrations Were Increased in Subjects with Metabolic Syndrome and Independently Associated with Fasting Plasma Glucose Levels. J Clin Med 2019;8

45. Fu Z, Gilbert ER, Liu D: Regulation of insulin synthesis and secretion and pancreatic Beta-cell dysfunction in diabetes. Curr Diabetes Rev 2013;9:25–53

46. Blum B, Hrvatin S, Schuetz C, Bonal C, Rezania A, Melton DA: Functional beta-cell maturation is marked by an increased glucose threshold and by expression of urocortin 3. Nat Biotechnol 2012;30:261–264

47. Sachdeva MM, Claiborn KC, Khoo C, Yang J, Groff DN, Mirmira RG, Stoffers DA: Pdx1 (MODY4) regulates pancreatic beta cell susceptibility to ER stress. Proc Natl Acad Sci U S A 2009;106:19090–19095

48. Adriaens C, Standaert L, Barra J, Latil M, Verfaillie A, Kalev P, Boeckx B, Wijnhoven PW, Radaelli E, Vermi W, Leucci E, Lapouge G, Beck B, van den Oord J, Nakagawa S, Hirose T, Sablina AA, Lambrechts D, Aerts S, Blanpain C, Marine JC: p53 induces formation of NEAT1 lncRNA-containing paraspeckles that modulate replication stress response and chemosensitivity. Nat Med 2016;22:861–868

49. Lepine S, Allegood JC, Park M, Dent P, Milstien S, Spiegel S: Sphingosine-1-phosphate phosphohydrolase-1 regulates ER stress-induced autophagy. Cell Death Differ 2011;18:350–361

50. Boyce M, Py BF, Ryazanov AG, Minden JS, Long K, Ma D, Yuan J: A pharmacoproteomic approach implicates eukaryotic elongation factor 2 kinase in ER stress-induced cell death. Cell Death Differ 2008;15:589–599

51. Kalsbeek A, la Fleur S, Fliers E: Circadian control of glucose metabolism. Mol Metab 2014;3:372–383

52. Perelis M, Marcheva B, Ramsey KM, Schipma MJ, Hutchison AL, Taguchi A, Peek CB, Hong H, Huang W, Omura C, Allred AL, Bradfield CA, Dinner AR, Barish GD, Bass J: Pancreatic beta cell enhancers regulate rhythmic transcription of genes controlling insulin secretion. Science 2015;350:aac4250

53. Allaman-Pillet N, Roduit R, Oberson A, Abdelli S, Ruiz J, Beckmann JS, Schorderet DF, Bonny C: Circadian regulation of islet genes involved in insulin production and secretion. Mol Cell Endocrinol 2004;226:59–66

54. Petrenko V, Gosmain Y, Dibner C: High-Resolution Recording of the Circadian Oscillator in Primary Mouse alpha- and beta-Cell Culture. Front Endocrinol (Lausanne) 2017;8:68

55. Groop L, Harno K: Diurnal pattern of plasma insulin and blood glucose during glibenclamide and glipizide therapy in elderly diabetics. Acta Endocrinol Suppl (Copenh) 1980;239:44–52

56. Polonsky KS, Given BD, Van Cauter E: Twenty-four-hour profiles and pulsatile patterns of insulin secretion in normal and obese subjects. J Clin Invest 1988;81:442–448

57. Katsuta H, Aguayo-Mazzucato C, Katsuta R, Akashi T, Hollister-Lock J, Sharma AJ, Bonner-Weir S, Weir GC: Subpopulations of GFP-marked mouse pancreatic beta-cells differ in size, granularity, and insulin secretion. Endocrinology 2012;153:5180–5187

58. Vafiadis P, Bennett ST, Colle E, Grabs R, Goodyer CG, Polychronakos C: Imprinted and genotype-specific expression of genes at the IDDM2 locus in pancreas and leucocytes. J Autoimmun 1996;9:397–403

59. Wang S, Flibotte S, Camunas-Soler J, MacDonald PE, Johnson JD: The type 1 diabetes at-risk allele at rs3842753 associates with increased beta cell INS mRNA in a meta-analysis of single cell RNA sequencing data. bioRxiv 2020:2020.2012.2006.413971

60. Yang BT, Dayeh TA, Kirkpatrick CL, Taneera J, Kumar R, Groop L, Wollheim CB, Nitert MD, Ling C: Insulin promoter DNA methylation correlates negatively with insulin gene expression and positively with HbA(1c) levels in human pancreatic islets. Diabetologia 2011;54:360–367

61. Nasteska D, Fine NHF, Ashford FB, Cuozzo F, Viloria K, Smith G, Dahir A, Dawson PWJ, Lai YC, Bastidas-Ponce A, Bakhti M, Rutter GA, Fiancette R, Nano R, Piemonti L, Lickert H, Zhou Q, Akerman I, Hodson DJ: PDX1(LOW) MAFA(LOW) beta-cells contribute to islet function and insulin release. Nat Commun 2021;12:674

62. Perl S, Kushner JA, Buchholz BA, Meeker AK, Stein GM, Hsieh M, Kirby M, Pechhold S, Liu EH, Harlan DM, Tisdale JF: Significant human beta-cell turnover is limited to the first three decades of life as determined by in vivo thymidine analog incorporation and radiocarbon dating. J Clin Endocrinol Metab 2010;95:E234–239

63. Teta M, Long SY, Wartschow LM, Rankin MM, Kushner JA: Very slow turnover of beta-cells in aged adult mice. Diabetes 2005;54:2557–2567

64. Veres A, Faust AL, Bushnell HL, Engquist EN, Kenty JH, Harb G, Poh YC, Sintov E, Gurtler M, Pagliuca FW, Peterson QP, Melton DA: Charting cellular identity during human in vitro beta-cell differentiation. Nature 2019;569:368–373

65. Thomsen SK, Raimondo A, Hastoy B, Sengupta S, Dai XQ, Bautista A, Censin J, Payne AJ, Umapathysivam MM, Spigelman AF, Barrett A, Groves CJ, Beer NL, Manning Fox JE, McCarthy MI, Clark A, Mahajan A, Rorsman P, MacDonald PE, Gloyn AL: Type 2 diabetes risk alleles in PAM impact insulin release from human pancreatic beta-cells. Nat Genet 2018;50:1122–1131

66. Nakayama M, Abiru N, Moriyama H, Babaya N, Liu E, Miao D, Yu L, Wegmann DR, Hutton JC, Elliott JF, Eisenbarth GS: Prime role for an insulin epitope in the development of type 1 diabetes in NOD mice. Nature 2005;435:220–223

67. Stadinski BD, Delong T, Reisdorph N, Reisdorph R, Powell RL, Armstrong M, Piganelli JD, Barbour G, Bradley B, Crawford F, Marrack P, Mahata SK, Kappler JW, Haskins K: Chromogranin A is an autoantigen in type 1 diabetes. Nat Immunol 2010;11:225–231

68. Han S, Donelan W, Wang H, Reeves W, Yang LJ: Novel autoantigens in type 1 diabetes. Am J Transl Res 2013;5:379–392

69. Rui J, Deng S, Arazi A, Perdigoto AL, Liu Z, Herold KC: beta Cells that Resist Immunological Attack Develop during Progression of Autoimmune Diabetes in NOD Mice. Cell Metab 2017;25:727–738

70. Dirice E, Kahraman S, De Jesus DF, El Ouaamari A, Basile G, Baker RL, Yigit B, Piehowski PD, Kim MJ, Dwyer AJ, Ng RWS, Schuster C, Vethe H, Martinov T, Ishikawa Y, Teo AKT, Smith RD, Hu J, Haskins K, Serwold T, Qian W-J, Fife BT, Kissler S, Kulkarni RN: Increased β-cell proliferation before immune cell invasion prevents progression of type 1 diabetes. Nature Metabolism 2019;

71. Raj A, Peskin CS, Tranchina D, Vargas DY, Tyagi S: Stochastic mRNA synthesis in mammalian cells. PLoS Biol 2006;4:e309

72. Vera M, Biswas J, Senecal A, Singer RH, Park HY: Single-Cell and Single-Molecule Analysis of Gene Expression Regulation. Annual review of genetics 2016;50:267–291

73. Suter DM, Molina N, Gatfield D, Schneider K, Schibler U, Naef F: Mammalian genes are transcribed with widely different bursting kinetics. Science 2011;332:472–474

74. Norris AJ, Stirland JA, McFerran DW, Seymour ZC, Spiller DG, Loudon AS, White MR, Davis JR: Dynamic patterns of growth hormone gene transcription in individual living pituitary cells. Mol Endocrinol 2003;17:193–202

75. Walsh HE, Shupnik MA: Proteasome regulation of dynamic transcription factor occupancy on the GnRH-stimulated luteinizing hormone beta-subunit promoter. Mol Endocrinol 2009;23:237–250

76. Harper CV, Featherstone K, Semprini S, Friedrichsen S, McNeilly J, Paszek P, Spiller DG, McNeilly AS, Mullins JJ, Davis JR, White MR: Dynamic organisation of prolactin gene expression in living pituitary tissue. J Cell Sci 2010;123:424–430

77. Phillips NE, Mandic A, Omidi S, Naef F, Suter DM: Memory and relatedness of transcriptional activity in mammalian cell lineages. Nat Commun 2019;10:1208

78. Nicolas D, Zoller B, Suter DM, Naef F: Modulation of transcriptional burst frequency by histone acetylation. Proc Natl Acad Sci U S A 2018;115:7153–7158

79. Nicolas D, Phillips NE, Naef F: What shapes eukaryotic transcriptional bursting? Mol Biosyst 2017;13:1280–1290

80. Atger F, Gobet C, Marquis J, Martin E, Wang J, Weger B, Lefebvre G, Descombes P, Naef F, Gachon F: Circadian and feeding rhythms differentially affect rhythmic mRNA transcription and translation in mouse liver. Proc Natl Acad Sci U S A 2015;112:E6579–6588

81. Molina N, Suter DM, Cannavo R, Zoller B, Gotic I, Naef F: Stimulus-induced modulation of transcriptional bursting in a single mammalian gene. Proc Natl Acad Sci U S A 2013;110:20563–20568

82. Pichon X, Lagha M, Mueller F, Bertrand E: A Growing Toolbox to Image Gene Expression in Single Cells: Sensitive Approaches for Demanding Challenges. Mol Cell 2018;71:468–480

83. Pichon X, Robert MC, Bertrand E, Singer RH, Tutucci E: New Generations of MS2 Variants and MCP Fusions to Detect Single mRNAs in Living Eukaryotic Cells. Methods Mol Biol 2020;2166:121–144

84. Vera M, Tutucci E, Singer RH: Imaging Single mRNA Molecules in Mammalian Cells Using an Optimized MS2-MCP System. Methods Mol Biol 2019;2038:3–20

85. Corish P, Tyler-Smith C: Attenuation of green fluorescent protein half-life in mammalian cells. Protein Eng 1999;12:1035–1040

86. Chowdhury A, Dyachok O, Tengholm A, Sandler S, Bergsten P: Functional differences between aggregated and dispersed insulin-producing cells. Diabetologia 2013;56:1557–1568

87. Scarl RT, Corbin KL, Vann NW, Smith HM, Satin LS, Sherman A, Nunemaker CS: Intact pancreatic islets and dispersed beta-cells both generate intracellular calcium oscillations but differ in their responsiveness to glucose. Cell Calcium 2019;83:102081

88. Kiepas A, Voorand E, Mubaid F, Siegel PM, Brown CM: Optimizing live-cell fluorescence imaging conditions to minimize phototoxicity. J Cell Sci 2020;133

89. Laissue PP, Alghamdi RA, Tomancak P, Reynaud EG, Shroff H: Assessing phototoxicity in live fluorescence imaging. Nat Methods 2017;14:657–661

90. Alvarez-Dominguez JR, Donaghey J, Rasouli N, Kenty JHR, Helman A, Charlton J, Straubhaar JR, Meissner A, Melton DA: Circadian Entrainment Triggers Maturation of Human In Vitro Islets. Cell Stem Cell 2020;26:108–122 e110

