## Supplemental figures for "Dynamic *Ins2* gene activity defines β-cell maturity states"

### Supplemental Figure 1

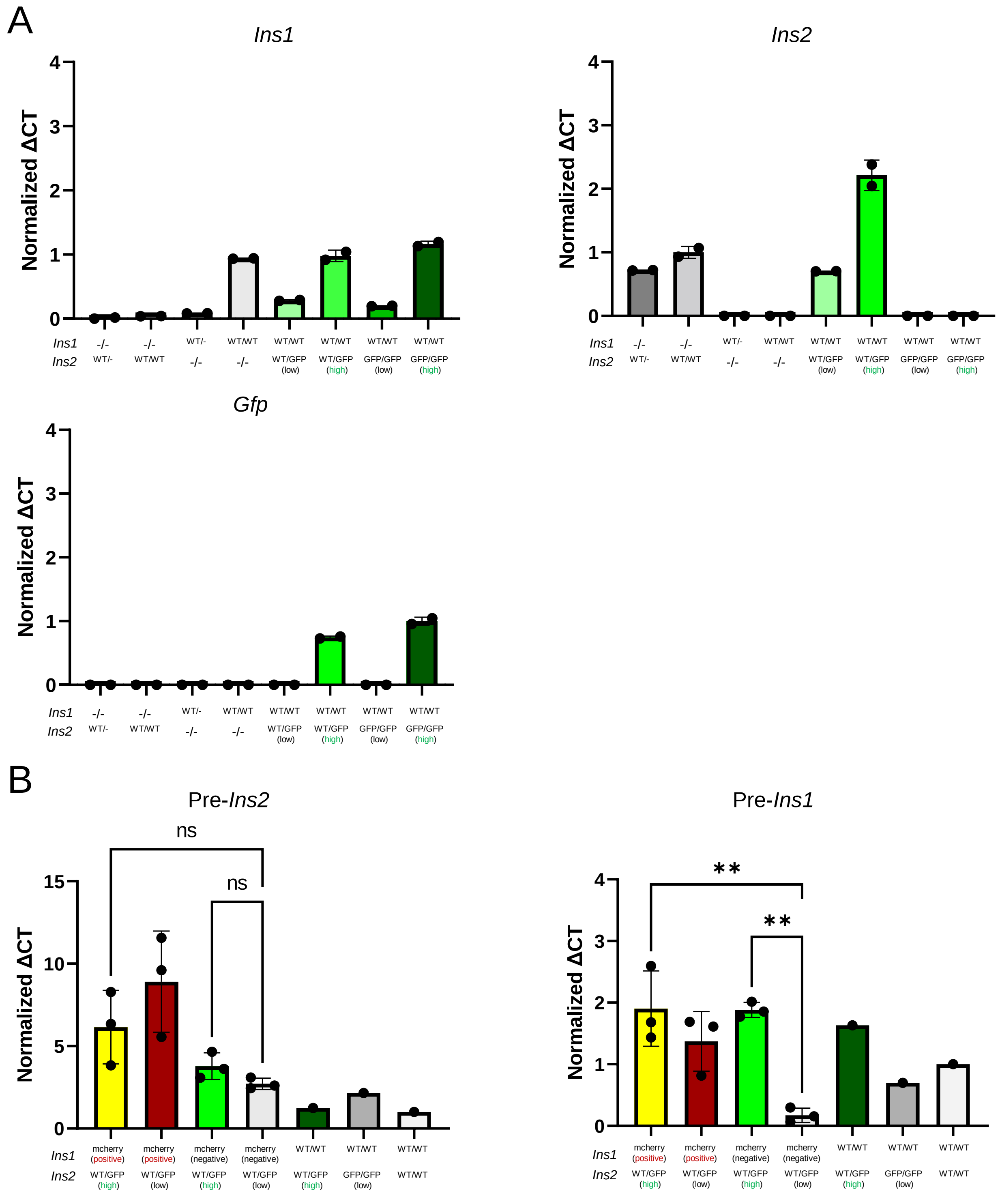


**Figure S1. qPCR results from *Ins2*$^{GFP}$ mice and *Ins2*$^{GFP/WT}$:*Ins1*-mCherry mice. (A)** mRNA of *Ins1*, *Ins2*, and *gfp* in various phenotypes of *Ins2*$^{GFP}$ mice. Units are relative expression calculated from ΔCT, then normalized to wild-type for *Ins1* and *Ins2*, and normalized to *Ins2*$^{GFP/GFP}$ mice for *gfp*. (n=1 mouse) **(B)** mRNA of pre-*Ins1* and pre-*Ins2* in *Ins2*$^{GFP/WT}$:*Ins1*-mCherry mice. Units are relative expression calculated from ΔCT, then normalized to wild-type (n=3 mice). One-way analysis of variance. * $p < 0.05$, ** $p < 0.01$, *** $p < 0.001$

### Supplemental Figure 2

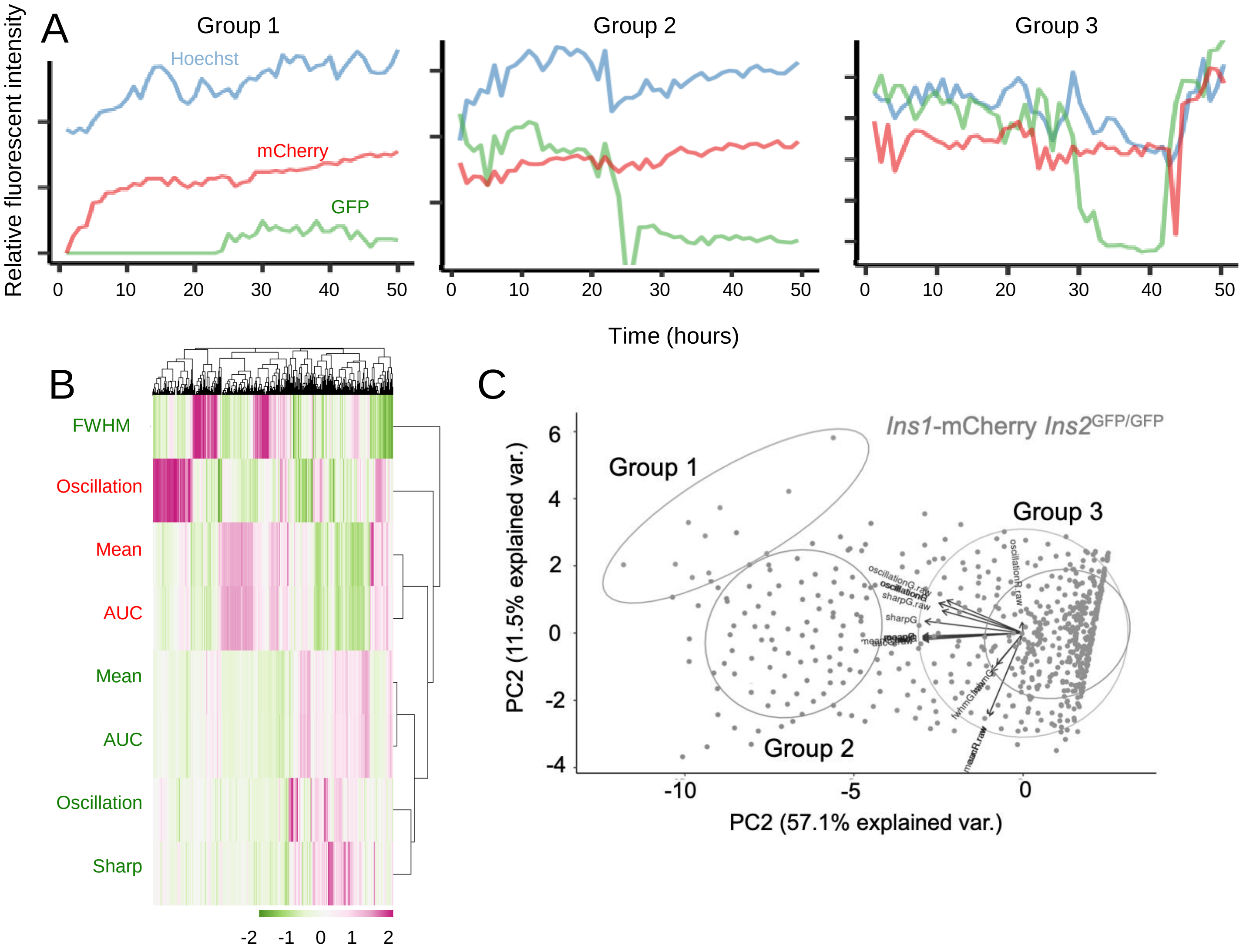


**Figure S2. Quantification of long term live cell imaging of *Ins2*$^{GFP/WT}$:*Ins1*-mCherry. (A)** Samples from three major groups identified from PCA analysis. **(B)** Heatmap with various calculations based on the red (mCherry) and green (GFP) wavelengths. Calculations include FWHM (full width at half maximum), Oscillation (mean average deviation), Mean, AUC (area under curve), and Sharp (number of sharp peaks over a cell's lifetime). **(C)** Principle component analysis based on calculations on the red and green wavelengths.

### Supplemental Figure 3

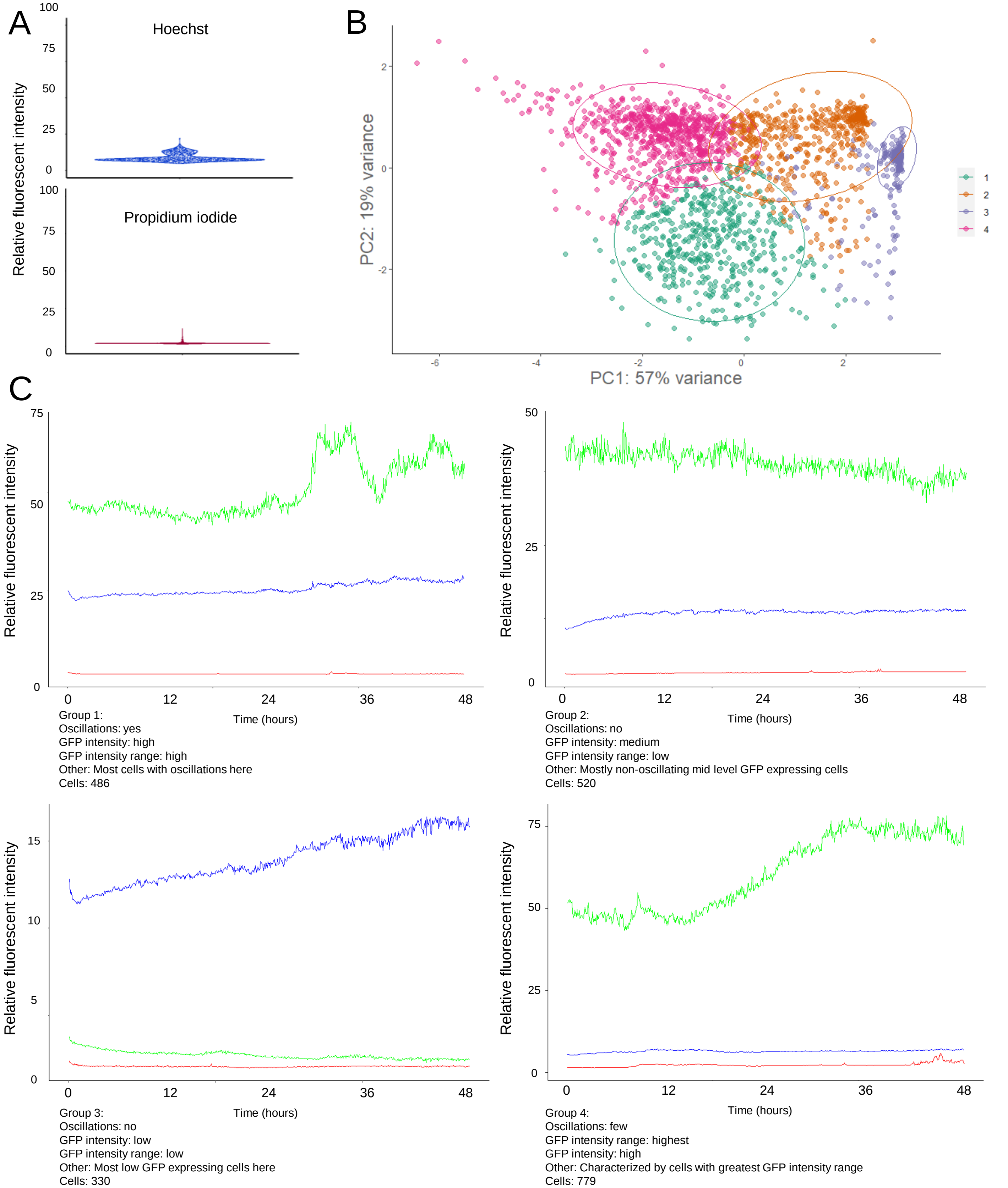


**Figure S3. Quantification of long term live cell imaging of *Ins2*$^{GFP/WT}$ cells. (A)** Lifetime fluorescent intensities of the blue and red wavelengths. **(B)** Principle component analysis on 2115 cells, based off cellular traits in the green wavelength (GFP). Principle components (eigenvectors) were calculated from calculations in Fig. 3E **(C)** Examples and observed traits from the four major populations of cells identified in PCA. Relative fluorescence intensity: fluorescence intensity relative to background, scaled 0-100.

### Supplemental Figure 4

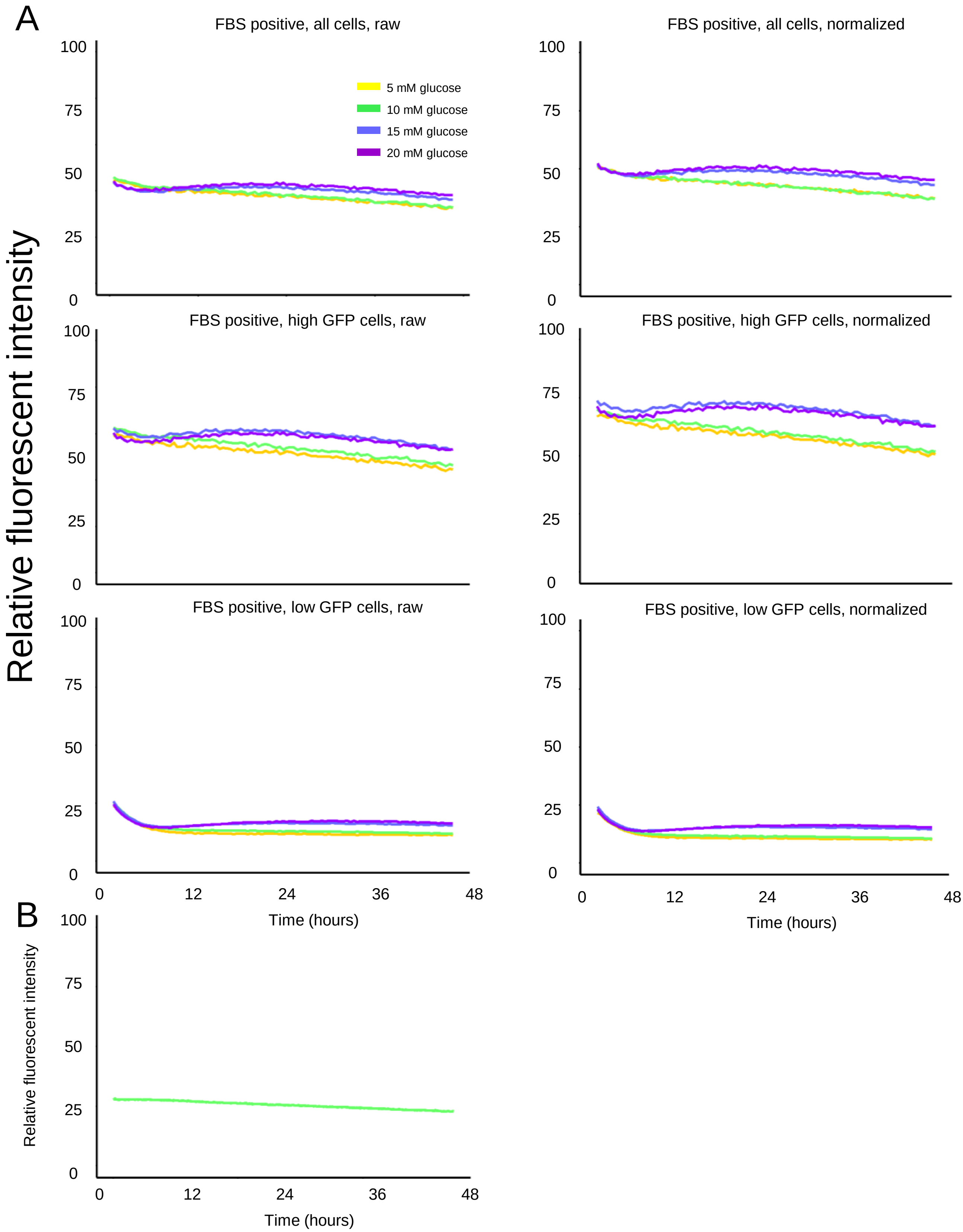


**Figure S4. GFP fluorescence over time. (A)** Average GFP fluorescent intensity over time of cells. Separated into all cells, high GFP expressing cells, and low GFP expressing cells and including raw and normalized data. Normalized data is normalized to the first two hours of imaging. **(B)** Experiment testing for effects of photo bleaching on GFP fluorescence over time. Relative fluorescent intensity: fluorescent intensity relative to background, scaled from 0-100.

### Supplemental Figure 5

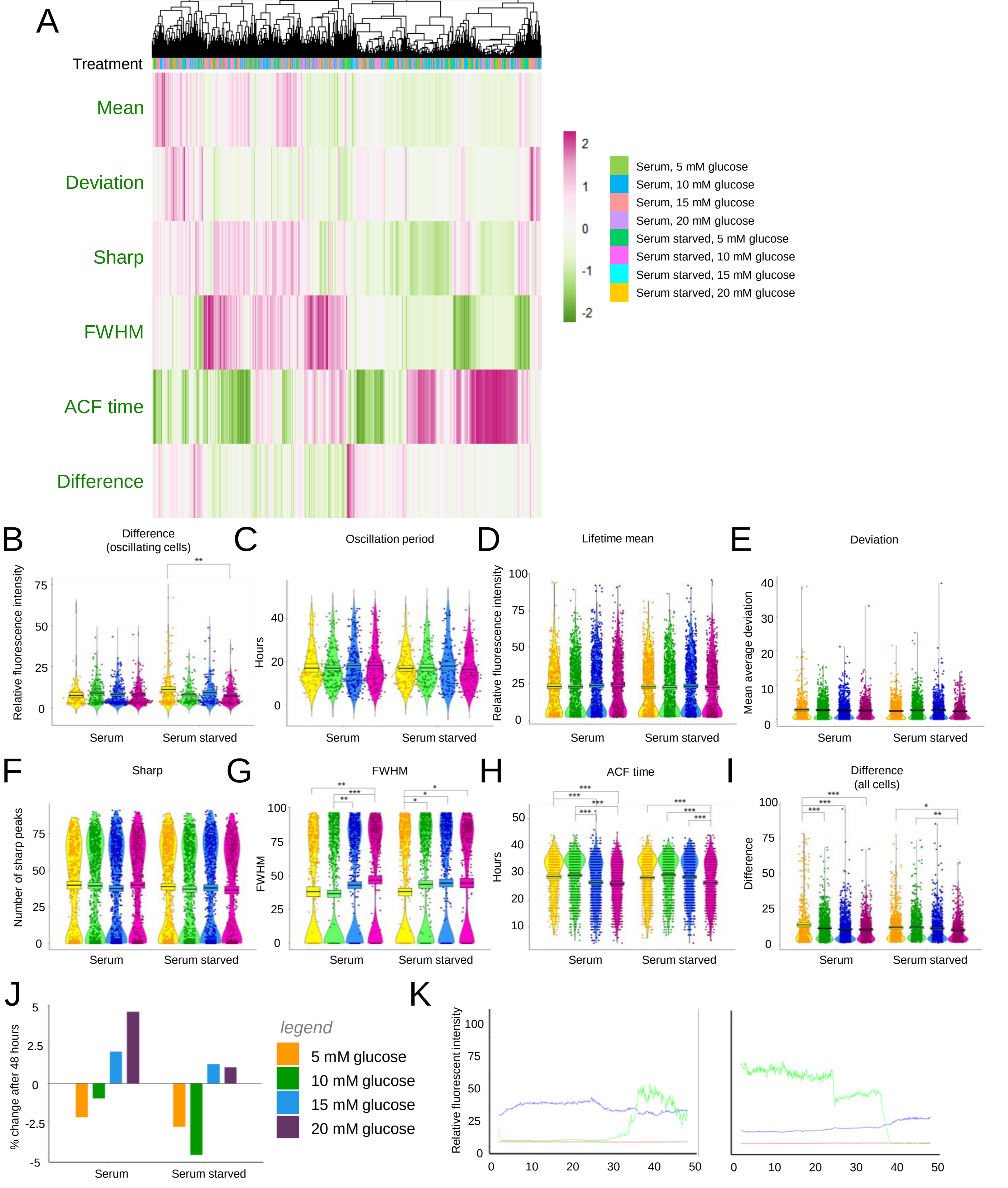


**Figure S5. Quantification of long term live cell imaging of *Ins2*$^{GFP/WT}$ cells in various culture conditions. (A)** Heatmap constructed with various calculations from the green (GFP) wavelength. **(B-I)** Visualization and statistical analyses on the various calculations from the green (GFP) wavelength. **(J)** Percent change in ratio of *Ins2*(GFP)$^{high}$ cells after 48 hours in various culture conditions. **(K)** Examples of cells transitioning to high and low GFP fluorescent states. Relative fluorescent intensity: fluorescent intensity relative to background, scaled from 0-100. Kruskal–Wallis one-way analysis of variance. * $p < 0.05$, ** $p < 0.01$, *** $p < 0.001$

### Supplemental Figure 6

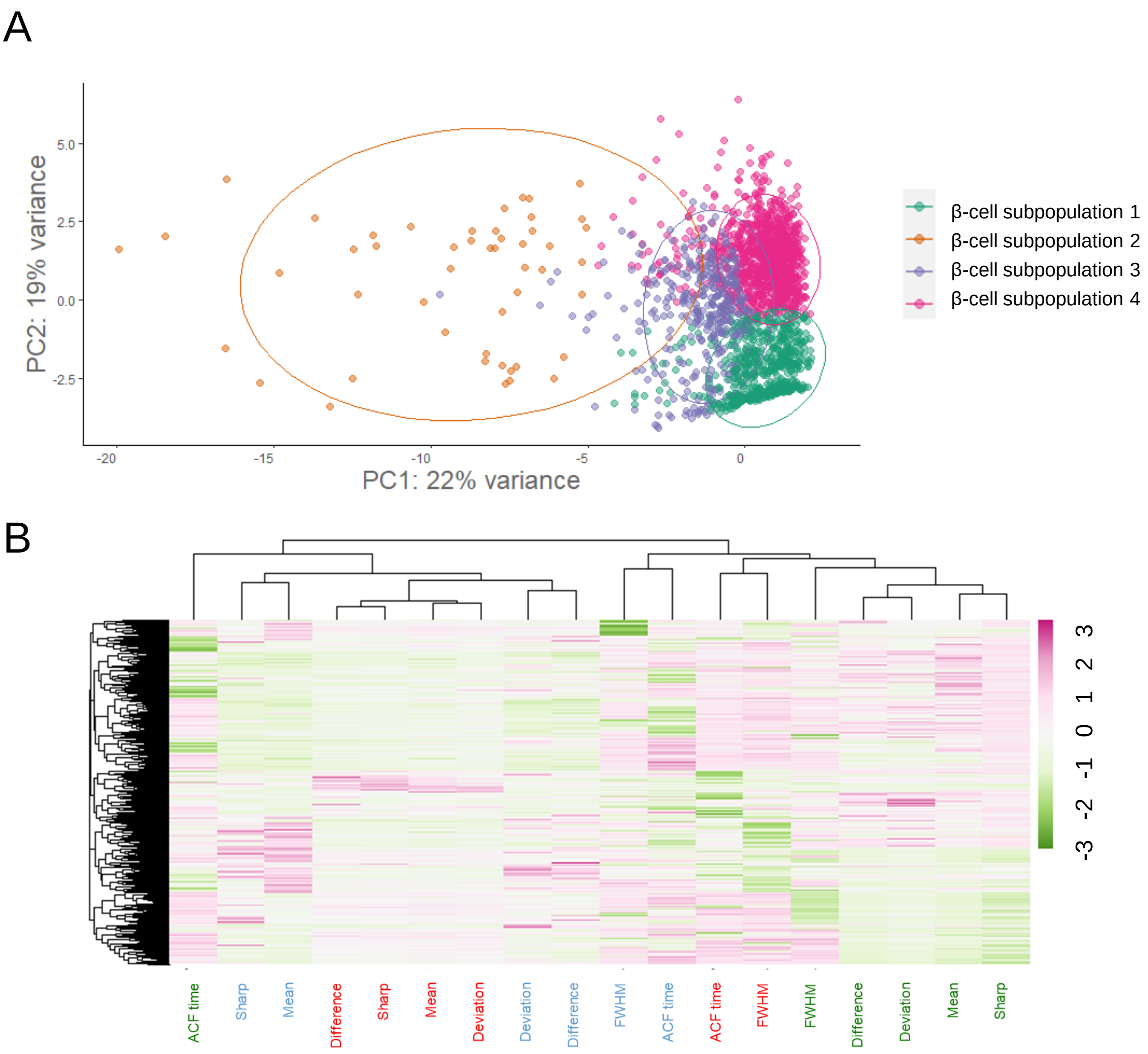


**Figure S6. Quantification of long term live cell imaging of Ins2$^{GFP/wt}$ cells in various culture conditions. (A)** Principle component analysis on cells in various culture conditions, with principle components (eigenvectors) based off cellular traits across all three wavelengths. **(B)** Heatmap constructed with various calculations from the blue (Hoechst cell nuclei dye), red (propidium iodide cell death marker) and green (GFP) wavelengths.

### Supplemental Figure 7

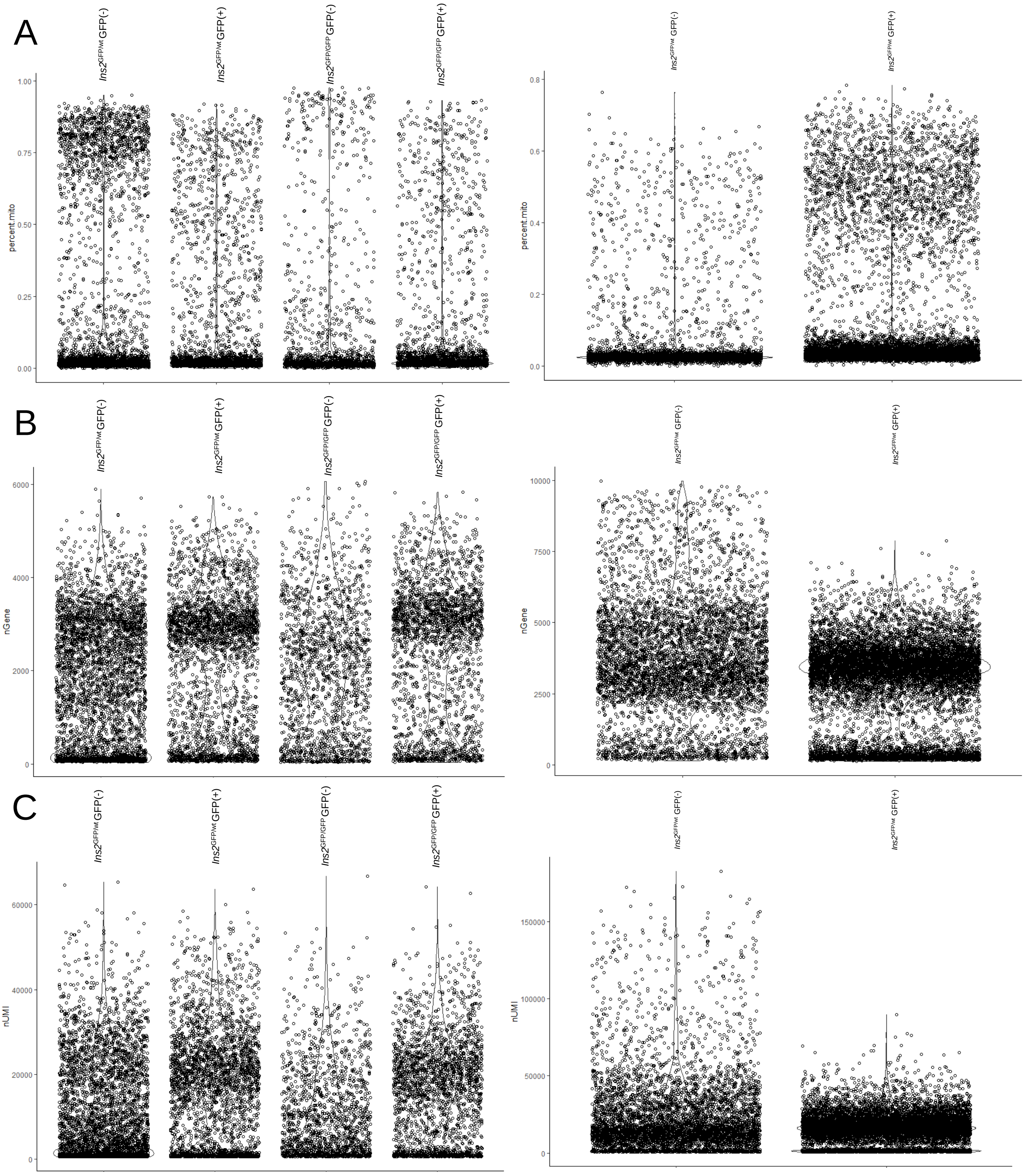


**Figure S7. Visualization of scRNA data prior to filtering.** Left: 60-week old mice. Right: 8-week old mice. **(A)** Percentage of mitochondria genes. **(B)** Number of genes per cell. **(C)** Number of genes per UMI.

### Supplemental Figure 8

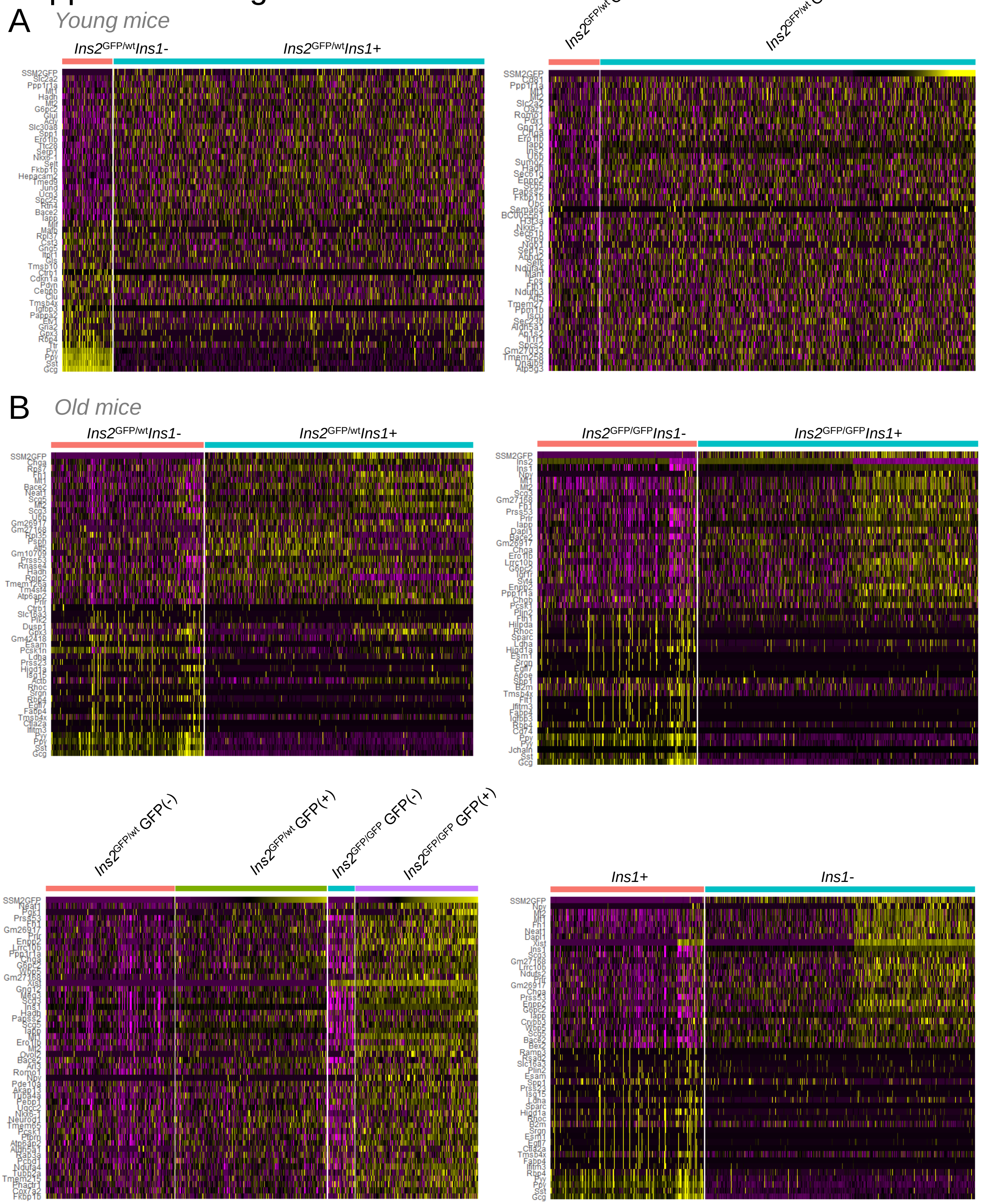


**Figure S8. Differential gene expression in *Ins2*(GFP)$^{LOW}$ and *Ins2*(GFP)$^{HIGH}$ β-cells.** Differential expression of *Ins1*+/GFP$^{LOW}$ vs *Ins1*+/GFP$^{HIGH}$ in **(A)** young *Ins2*$^{WT/GFP}$ **(B)** Old *Ins2*$^{GFP/WT}$, *Ins2*$^{GFP/GFP}$ and pooled genotypes. Genes are selected with 0.25 $\log_2$FC and expressed in at least 10% of cells in either group.

### Supplemental Figure 9

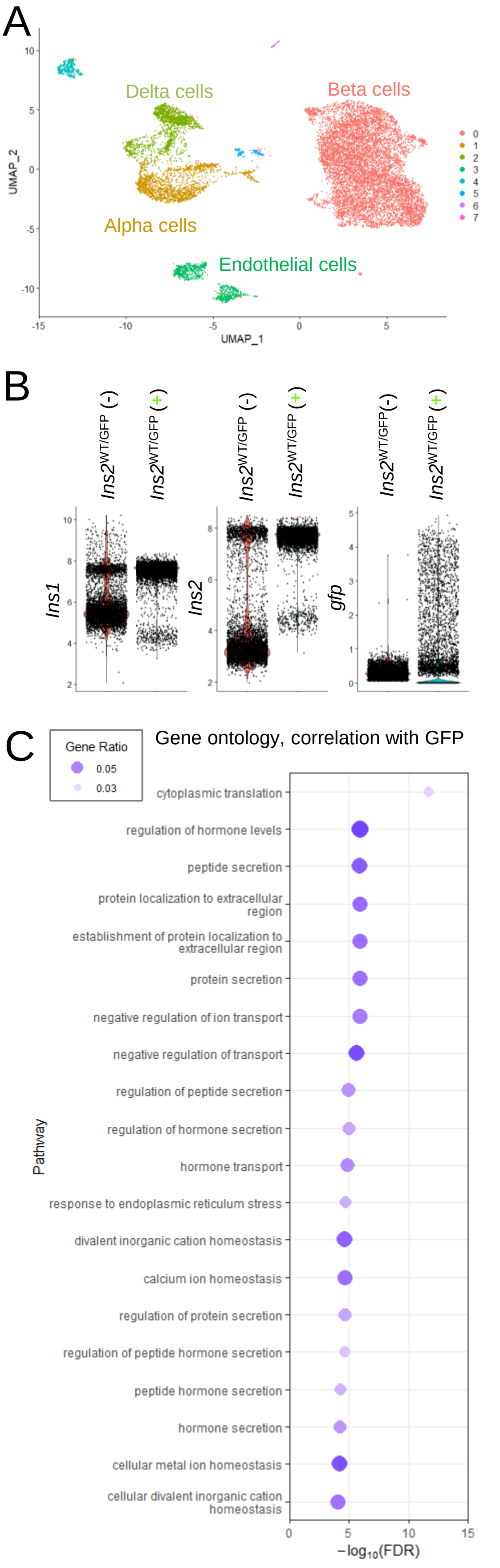


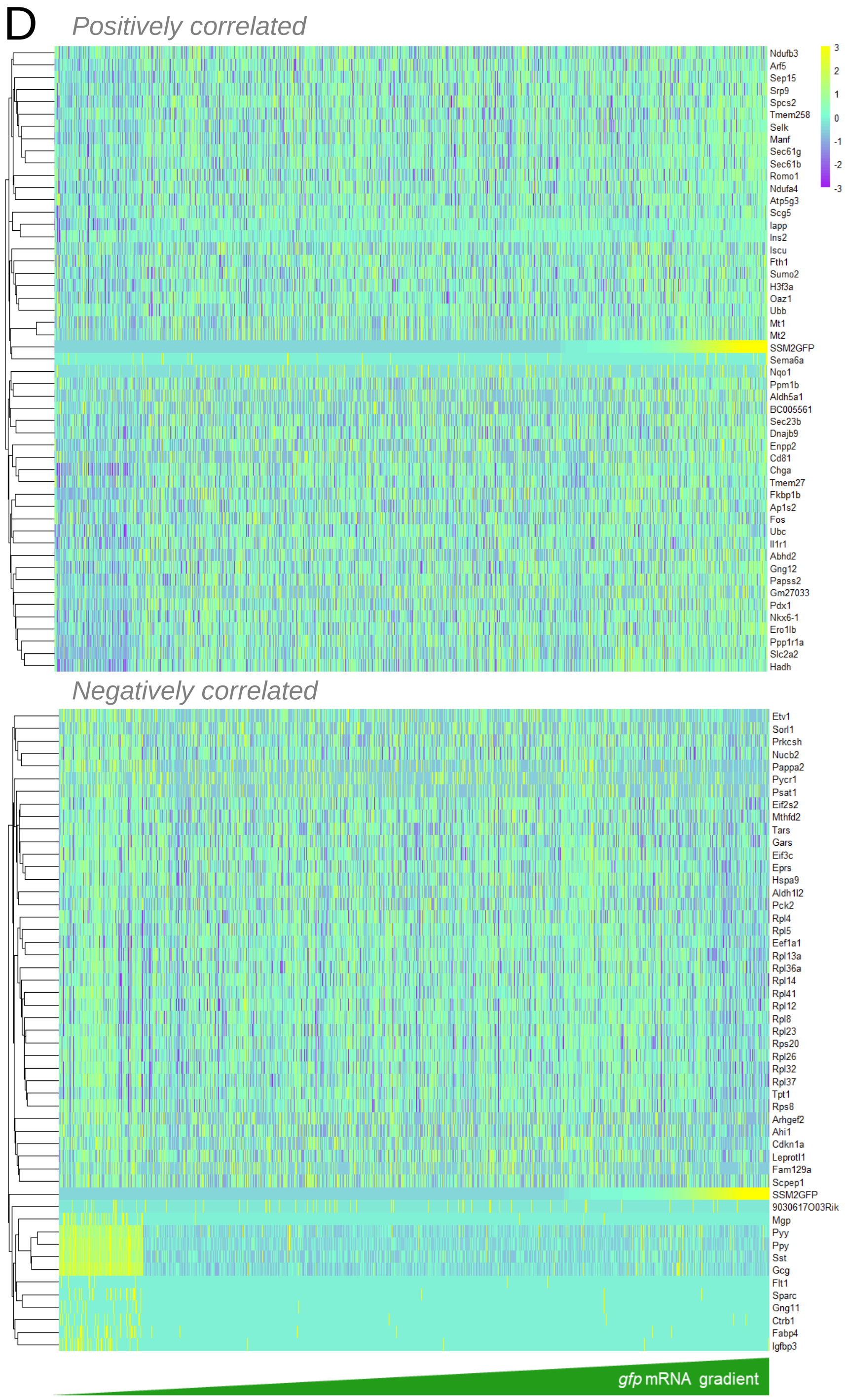


**Figure S9. Single cell RNA sequencing analysis of *Ins2*(GFP)$^{LOW}$ and *Ins2*(GFP)$^{HIGH}$ β-cells in young *Ins2*$^{WT/GFP}$ heterozygous mice. (A)** UMAP plot of all cells isolated and dispersed from *Ins2*$^{WT/GFP}$ mice and FACS purified into either GFP-negative (-) or GFP-positive (+) groups. **(B)** Log-normalized counts of *Ins1, Ins2,* and *gfp* mRNA quantification distributions from GFP-negative or GFP-positive single β-cells. **(C)** Gene Ontology Categories that are driven by genes differentially expressed in *Ins2*(GFP)$^{LOW}$ versus *Ins2*(GFP)$^{HIGH}$ β-cells. **(D)** Individual genes that are differentially expressed as a function of *gfp* mRNA expression.

### Supplemental Figure 10

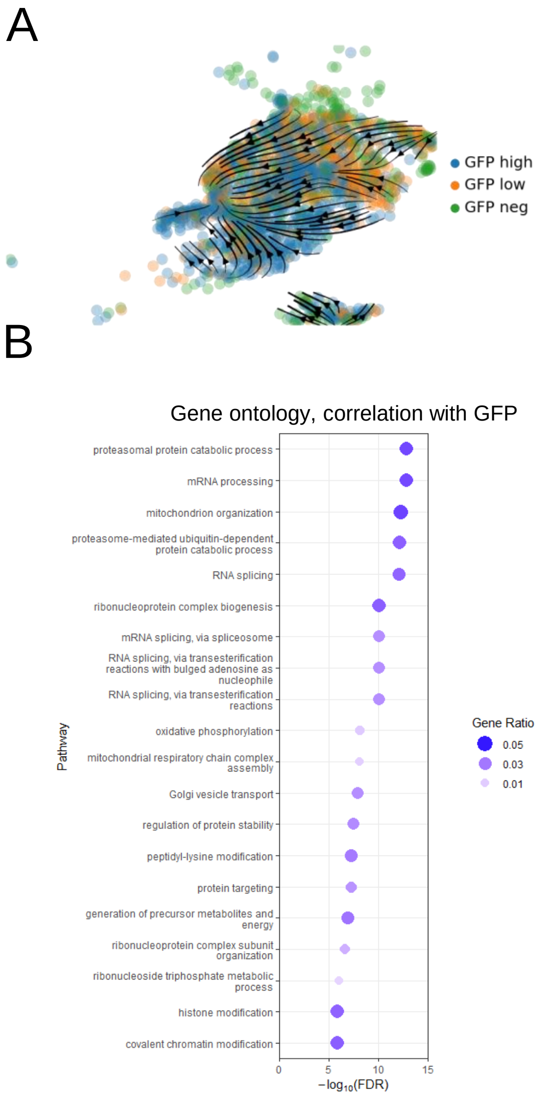


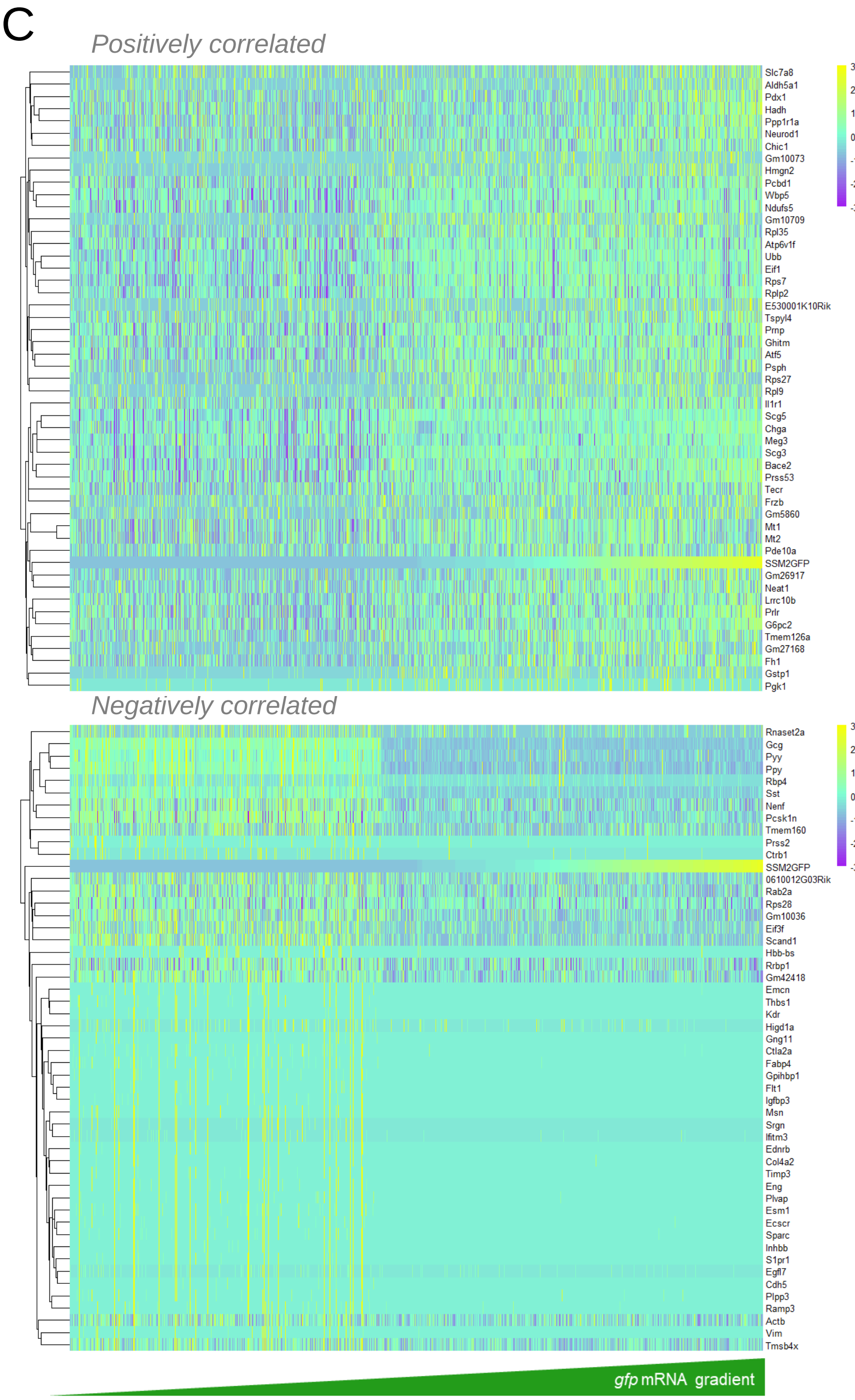


**Figure S10. Single cell RNA sequencing analysis of *Ins2*(GFP)$^{LOW}$ and *Ins2*(GFP)$^{HIGH}$ β-cells in old *Ins2*$^{GFP/GFP}$ homozygous mice. (A)** UMAP projection of β-cells isolated and dispersed from *Ins2*$^{GFP/GFP}$ homozygous mice with RNA velocity vectors overlaid. **(B)** Gene Ontology Categories that are driven by genes differentially expressed in *Ins2*(GFP)$^{LOW}$ versus *Ins2*(GFP)$^{HIGH}$ β-cells. **(C)** Individual genes that are differentially expressed as a function of *gfp* mRNA expression.

### Supplemental Figure 11

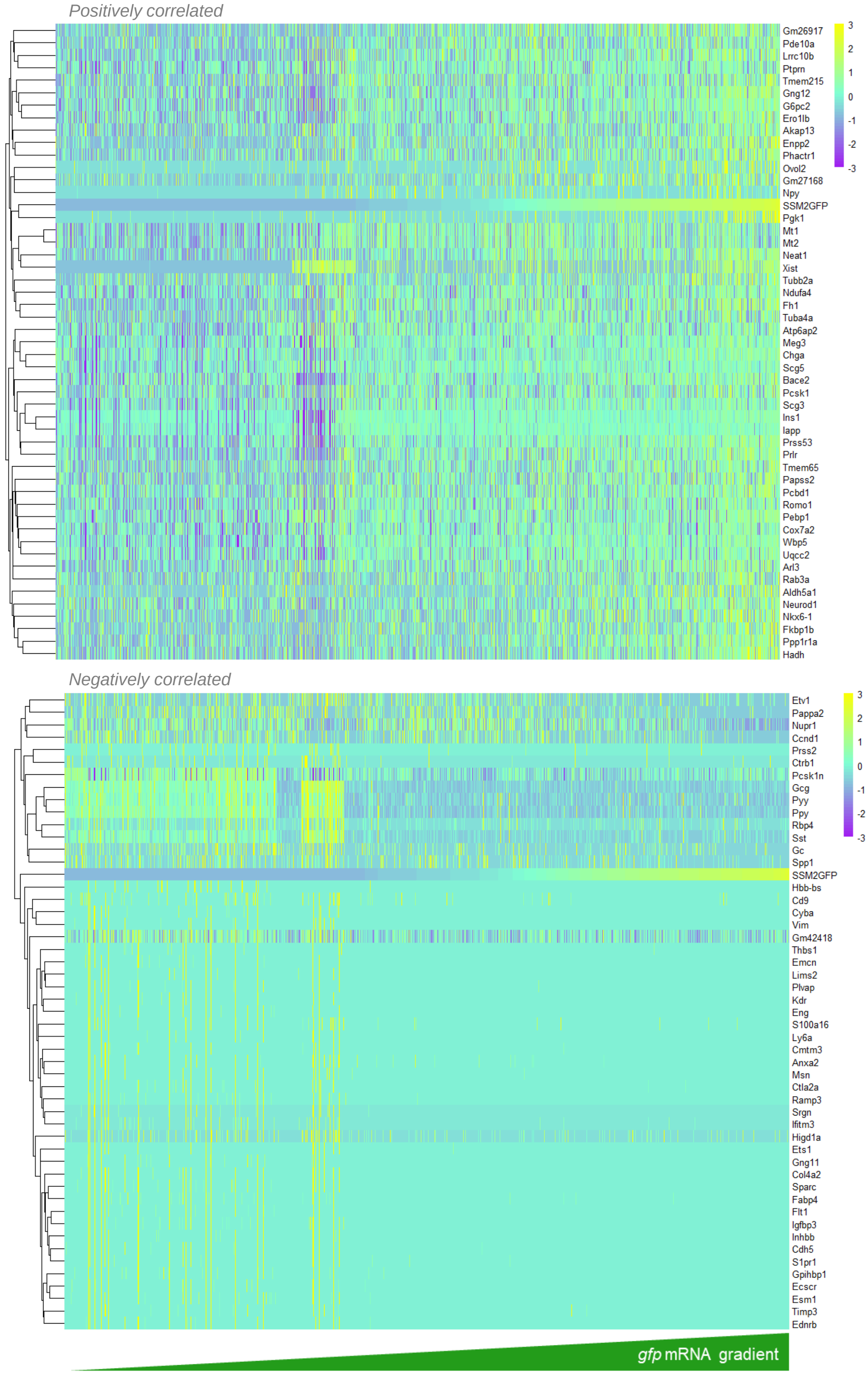


**Figure S11. Differential gene expression according to *gfp* mRNA levels in the combined samples of old *Ins2*$^{GFP/WT}$ heterozygous and *Ins2*$^{GFP/GFP}$ homozygous mice.** Single-cell RNA sequencing in *Ins1* expressing β-cells from *Ins2*$^{GFP}$ knock-in mice showing differentially expressed genes in heatmap form.

### Supplemental Figure 12

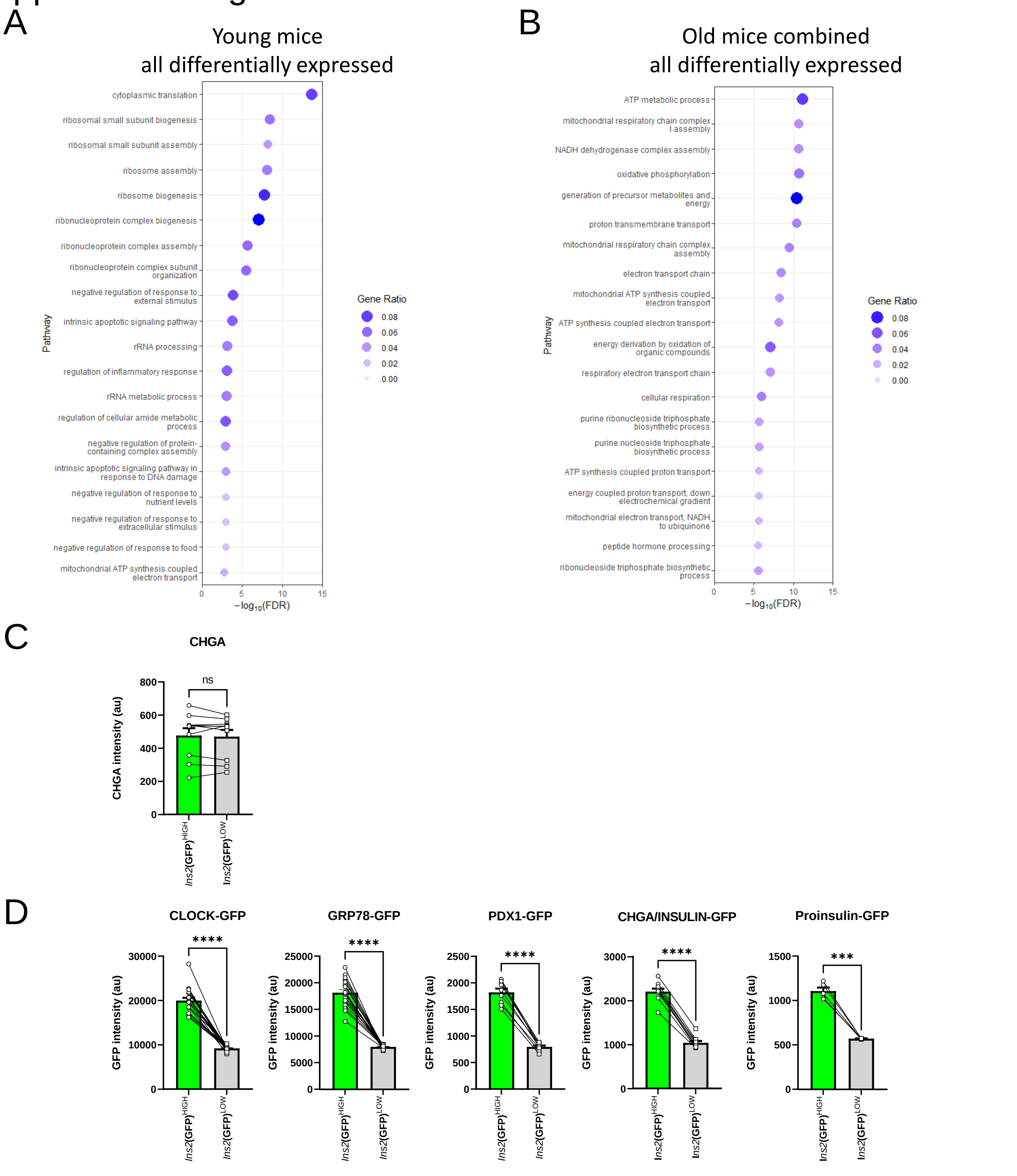


**Figure S12. Pathway enrichment analysis of differentially expressed genes in the combined samples of young and old *Ins2*$^{GFP}$ knock-in mice. (A)** Single-cell RNA sequencing in *Ins1* expressing β-cells from young 8-week old *Ins2*$^{GFP}$ knock-in mice showing differentially expressed pathways. **(B)** Single-cell RNA sequencing in *Ins1* expressing β-cells from combined samples of old *Ins2*$^{GFP/WT}$ heterozygous and *Ins2*$^{GFP/GFP}$ homozygous mice and knock-in mice showing differentially expressed pathways. **(C)** Quantification of staining for CHGA protein in pancreas sections from *Ins2*$^{WT/GFP}$ heterozygous mice. **(D)** Quantification of GFP intensity, comparing high GFP expressing regions and low GFP expressing regions, used in immunofluorescent staining for proteins of interest. Paired t-test. * $p < 0.05$, ** $p < 0.01$, *** $p < 0.001$, **** $p < 0.0001$.
